# Human marginal zone B cell development from early T2 progenitors

**DOI:** 10.1101/2020.09.24.311498

**Authors:** Thomas J. Tull, Michael J. Pitcher, William Guesdon, Jacqueline H. Siu, Cristina Lebrero-Fernández, Yuan Zhao, Nedyalko Petrov, Susanne Heck, Richard Ellis, Pawan Dhami, Ulrich D. Kadolsky, Michelle Kleeman, Yogesh Kamra, David J. Fear, Susan John, Wayel Jassem, Richard W. Groves, Jeremy D. Sanderson, Michael D. Robson, David D’Cruz, Mats Bemark, Jo Spencer

## Abstract

B cells emerge from the bone marrow as transitional (TS) B cells that differentiate through T1, T2 and T3 stages to become naïve B cells. We have identified a bifurcation of human B cell maturation from the T1 stage forming IgM^hi^ and IgM^lo^ developmental trajectories. IgM^hi^ T2 cells have higher expression of α4β7 integrin and lower expression of IL4 receptor (IL4R) compared to the IgM^lo^ branch and are selectively recruited into gut-associated lymphoid tissue. IgM^hi^ T2 cells also share transcriptomic features with marginal zone B cells (MZB). Lineage progression from T1 cells to MZB via an IgM^hi^ trajectory is identified by pseudotime analysis of scRNA-sequencing data. Reduced frequency of IgM^hi^ gut homing T2 cells is observed in severe SLE and is associated with reduction of MZB and their putative IgM^hi^ precursors. The collapse of the gut-associated MZB maturational axis in severe SLE affirms its existence and importance for maintaining health.

## Introduction

Transitional (TS) B cells are the immature B cells in human blood from which all mature B cells develop. Following emigration from bone marrow, TS B cells mature through T1, T2 and T3 phases, when autoreactive cells are depleted (Palanichamy et al., 2009; Suryani et al., 2010; Yurasov et al., 2005).

In mice a B cell lineage split that is dependent on B cell receptor (BCR) engagement and the serine/threonine kinase Taok3 is initiated at the T1 phase (Hammad et al., 2017). This directs B cells towards marginal zone B (MZB) cell fate, requiring subsequent Notch2 cleavage by a disintegrin and metalloproteinase-containing protein-10 (ADAM10).

MZB lineage progression in humans is not clearly understood, or indeed, universally accepted. A MZB precursor (MZP) population has been proposed that undergoes terminal differentiation to MZB following NOTCH2 ligation and can be discriminated from naïve B cells by expression of high levels of IgM (IgM^hi^), CD24 and the glycosylation-dependent epitope CD45RB^MEM55^ (referred to here as CD45RB). An additional CD45RB^hi^ IgM^hi^ population that lacks the ABCB1 cotransporter has previously been referred to as T3’, although the relationship between this subset, MZB and MZP is unclear (Bemark et al., 2013; Descatoire et al., 2014; Koethe et al., 2011; Zhao et al., 2018).

In humans, MZB develop over the first 2 years of life and are important for immunity against encapsulated bacteria (Weller et al., 2004). They undergo a phase of clonal expansion and receptor diversification in the germinal centres (GC) of gut-associated lymphoid tissue (GALT) (Zhao et al., 2018) (Weill and Reynaud, 2019). The shared expression of MAdCAM1 between the splenic marginal zone reticular cells and GALT high endothelial venules (HEV) creates the potential to recruit B cells to both sites mediated by α4β7 integrin binding (Kraal et al., 1995; Vossenkamper et al., 2013). We have described the expression of β7 integrin (used here and previously as a surrogate for α4β7) by T2 B cells in humans and observed their selective recruitment into GALT where they become activated (Vossenkamper et al., 2013). Therefore, exposure to the GALT microenvironment could be associated with multiple stages of MZB cell development from as early as the T2 stage.

The systemic autoimmune disease systemic lupus erythematosus (SLE), in particular the severe variant lupus nephritis (LN), has markedly distorted profiles of B cell subsets in blood. The TS pool is expanded, as is the B cell subset lacking both CD27 and IgD (so called ‘double negative’ or DN B cells) (Landolt-Marticorena et al., 2011; Wei et al., 2007).

Disproportionate expansion of a population of DN cells lacking expression of CD21 and CXCR5 and with upregulated CD11c (DN2 cells) is a particular feature of LN (Jenks et al., 2018). DN2 cells may be derived from activated naïve B cells (aNAV), driven by TLR7 engagement, resulting in the generation of self-reactive antibody producing plasma cells (Jenks et al., 2018; Tipton et al., 2015). Interestingly, a recent study of a cohort of newly diagnosed patients with SLE demonstrated that MZB may be reduced in frequency (Zhu et al., 2018). Since we have previously shown that TS B cells in SLE may have significantly reduced expression of β7 integrin, we were interested to know if this may be associated with defective MZB development and the increase in aNAV and DN2 cells.

Here, we identify bifurcation in human B cell development from the T2 stage. Cells in one branch are IgM^hi^, express β7 integrin and are gut homing. Cells in the alternative IgM^lo^ branch have high expression of IL4R, lower expression of β7 integrin and do not tend to enter the gut. Transcriptomically, IgM^hi^ T2 cells share features with MZB. B cell development progresses from T1 to MZB via an IgM^hi^ trajectory by pseudotime analysis. IgM^hi^ T2 cells are stably IgM^hi^ in culture and have a greater tendency to make IL10 than IgM^lo^ cells. Markedly reduced frequency of IgM^hi^β7^hi^ T2 cells was seen in patients with severe SLE and this was associated with stark reduction in cell populations associated with MZB development. Our data link reduced access of IgM^hi^ T2 cells to GALT with defects in all stages of MZB differentiation and enables the assimilation of these elements of human MZB differentiation into a model of human B cell development.

## Results

### Segregation of B cell phenotypes from T2 through naïve B cell subsets

In mice, B cells commit to MZB differentiation soon after bone marrow emigration at the T1 stage. To seek evidence of this in humans, a deep phenotypic analysis of peripheral blood mononuclear cells (PBMC) from healthy control donors (HCD) was undertaken by mass cytometry (Fig. S1 A, B and C). SPADE on viSNE identified B cell subsets including TS B cells represented by CD27^-^IgD^+^CD24^+++/++^CD38^+++/++^ nodes that included CD10^+^ T1 and T2 cells as well as CD10^-^ T3 cells (Fig. 1 A)(Qiu et al., 2011; Zhao et al., 2018). T3 cells can only be definitively distinguished from naïve cells by their failure to extrude dyes such as rhodamine 123 (R123) due to lack of the ABCB1 cotransporter (Wirths and Lanzavecchia, 2005) (Fig. S1 D). Since mass cytometry cannot be used to detect dye extrusion, the boundary between T3 and naïve B cells was estimated to generate the TS SPADE bubble. As previously reported, CD27^+^ B cells included CD27^hi^ and CD27^lo^ cells (Fig. 1 A) (Grimsholm et al., 2020).

**Figure 1.**
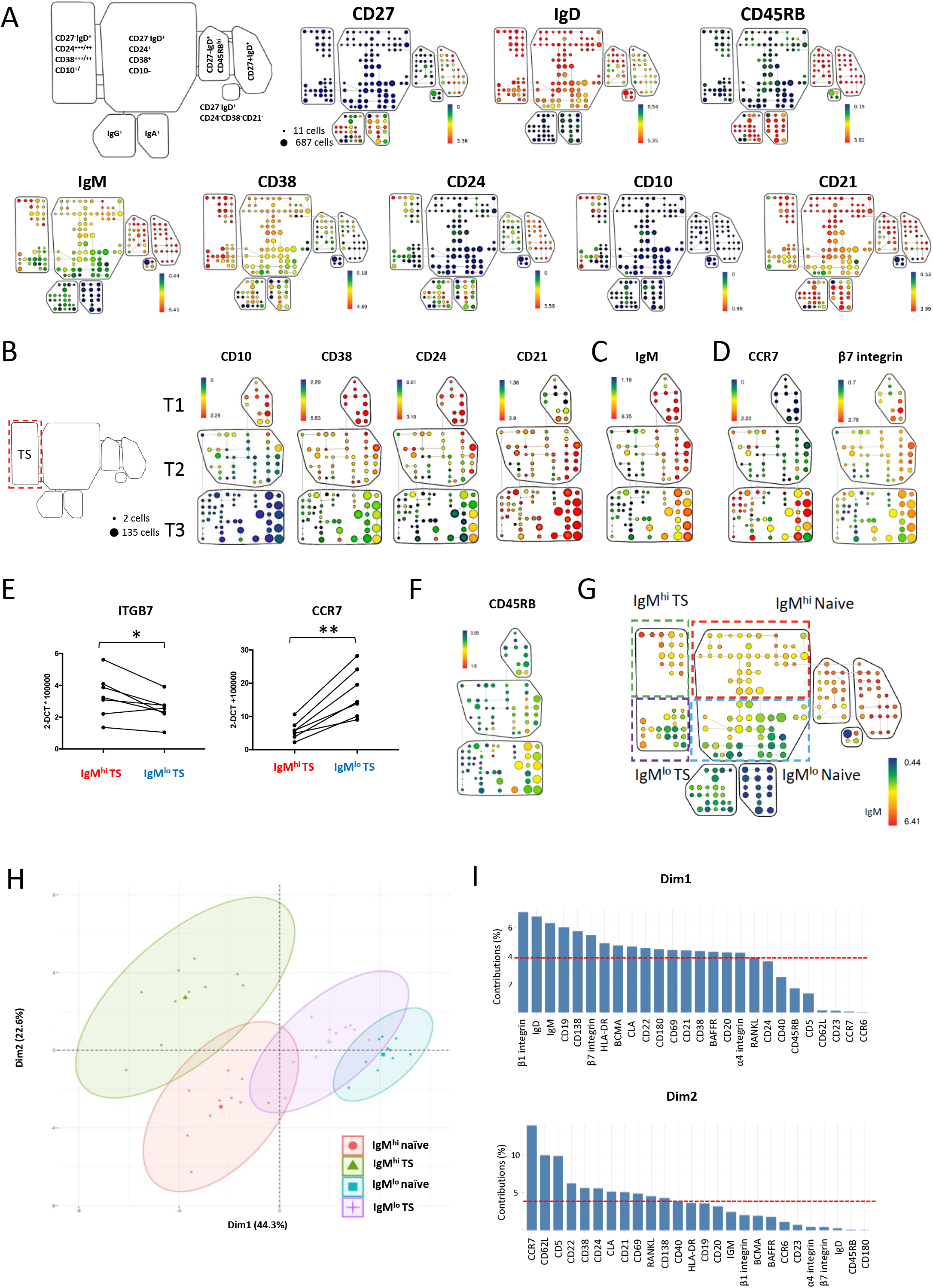
Segregation of B cell phenotypes from T2 through naïve B cell subsets. A) SPADE on viSNE plots generated using the following markers; CD10, CD24, CD38, CD27, CD45RB IgD, IgM, IgA and IgG. The plots are from a representative female HCD and depict expression of lineage markers used to identify B cell subsets in HCD PBMC. B) SPADE on viSNE plots of TS B cells exported from the TS SPADE bubble in Figure part A and generated by re-running viSNE using all expressed B cell markers. The SPADE plots depicted are from a representative female HCD. TS B cell populations were defined as T1 (CD10^+^CD24^+++^CD38^+++^CD21^lo^), T2 (CD10^+^CD24^++^CD38^++^CD21^hi^) and T3 (CD10^-^ CD24^+^CD38^+^CD21^hi^). C) SPADE trees demonstrating that T2 and T3 TS B cells have prominent IgM^hi^ and IgM^lo^ subpopulations. D) SPADE trees demonstrating that IgM^hi^ T2 B cells have higher expression of β7 integrin and lower expression of CCR7 than IgM^lo^ T2 B cells. E) qPCR validation of CCR7 and ITGB7 (β7 integrin) from sorted subsets expressed as ΔCT values relative to 18S endogenous control (paired t test). F) SPADE aligns IgM^hi^ T2 cells with IgM^hi^ T3 cells with relatively high expression of CD45RB. G) A SPADE on viSNE plot from Figure part A demonstrating the identification of IgM^hi^ TS and naïve B cell populations. H) A principle component analysis plot generated using all expressed markers on IgM^hi^ and IgM^lo^ subsets identified in Figure part G. Data points represent individual donors and are surrounded by 95% confidence ellipses with a larger central mean data marker. I) Variable contribution bar graphs demonstrate that homing receptors are major contributors to PCA1 and PCA2 in Figure part G. The dashed red reference line represents the value were the contribution uniform.

To perform a deep phenotypic analysis of TS B cells, events within the TS bubble identified in Fig. 1 A were exported and re-clustered by SPADE on viSNE using all expressed panel markers and then grouped according to gradients of loss of CD10, CD38 and CD24 and gain of CD21 corresponding to T1, T2 and T3 stages of differentiation (Fig. 1 B)(Bemark, 2015). The SPADE trees branched, forming 2 chains of nodes that each extended through the T2 and T3 SPADE bubbles with no lateral connections between the branches. Branches differed most notably in their expression of IgM (Fig. 1 C). IgM^hi^ T2 B cells also had lower expression of CCR7 but higher expression of β7 integrin than IgM^lo^ T2 cells by mass cytometry. This was validated by qPCR (Fig. 1 D and E). In addition, IgM^hi^ T3 cells had higher median expression of CD24 and CD45RB than IgM^lo^ T3 cells (Fig. 1 B and F).

In the SPADE analysis of all CD19^+^ B cells, nodes representing IgM^hi^ TS B cells were continuous with IgM^hi^ naïve B cells and IgM^lo^ TS B cells were continuous with IgM^lo^ naïve cells (Fig. 1 A and G). Principle component analysis (PCA) using all markers expressed by cells in nodes identified in Fig. 1 G grouped IgM^hi^ TS B cells closest to IgM^hi^ naïve B cells and most distant to IgM^lo^ TS B cells (Fig. 1 H). The major contributors to PCA1 and PCA2 in addition to IgM and IgD were mediators of cell traffic (Fig. 1 I).

Human B cells therefore segregate phenotypically as T1 cells enter into the T2 stage, forming two branches that differ in their expression of IgM and in markers of migratory potential. IgM^hi^ T2 cells resemble IgM^hi^ naïve cells more closely than they resemble IgM^lo^ T2 cells with which they share markers of differentiation.

### Gut-associated lymphoid tissue is enriched in IgM^hi^ T2 cells

Human TS B cells can home to GALT where they become activated (Vossenkamper et al., 2013). To determine whether the high expression of β7 integrin on IgM^hi^ TS B cells is associated with selective recruitment into GALT, mass cytometry was used to compare B cells isolated from paired blood and gut biopsies from individuals undergoing surveillance colonoscopies (n=7) (Fig. S1 E, F and G). TS B cells are a small subset and due to low mononuclear cell yields from GALT biopsies, data from individual samples were concatenated. SPADE on viSNE was then used to identify CD10^+^ T1 and T2 B cells within the total CD19^+^ population (Fig. 2 A and S1 H). The undirected clustering algorithm FlowSOM (Van Gassen et al., 2015) was then used to group T1 and T2 B cells. This identified six metaclusters and the identity of each cluster was deduced from the relative expression of CD21, CD24, CD38 and IgM (Fig. 2 B and C). This demonstrated that IgM^hi^ T2 cells are enriched in GALT, whereas both T1 and IgM^lo^ T2 cells are depleted compared to PBMC (Fig. 2 D and E). Within GALT IgM^hi^ T2 cells had higher expression of the activation markers CD69 and CD80 than PBMC (Fig. 2 E).

**Figure 2.**
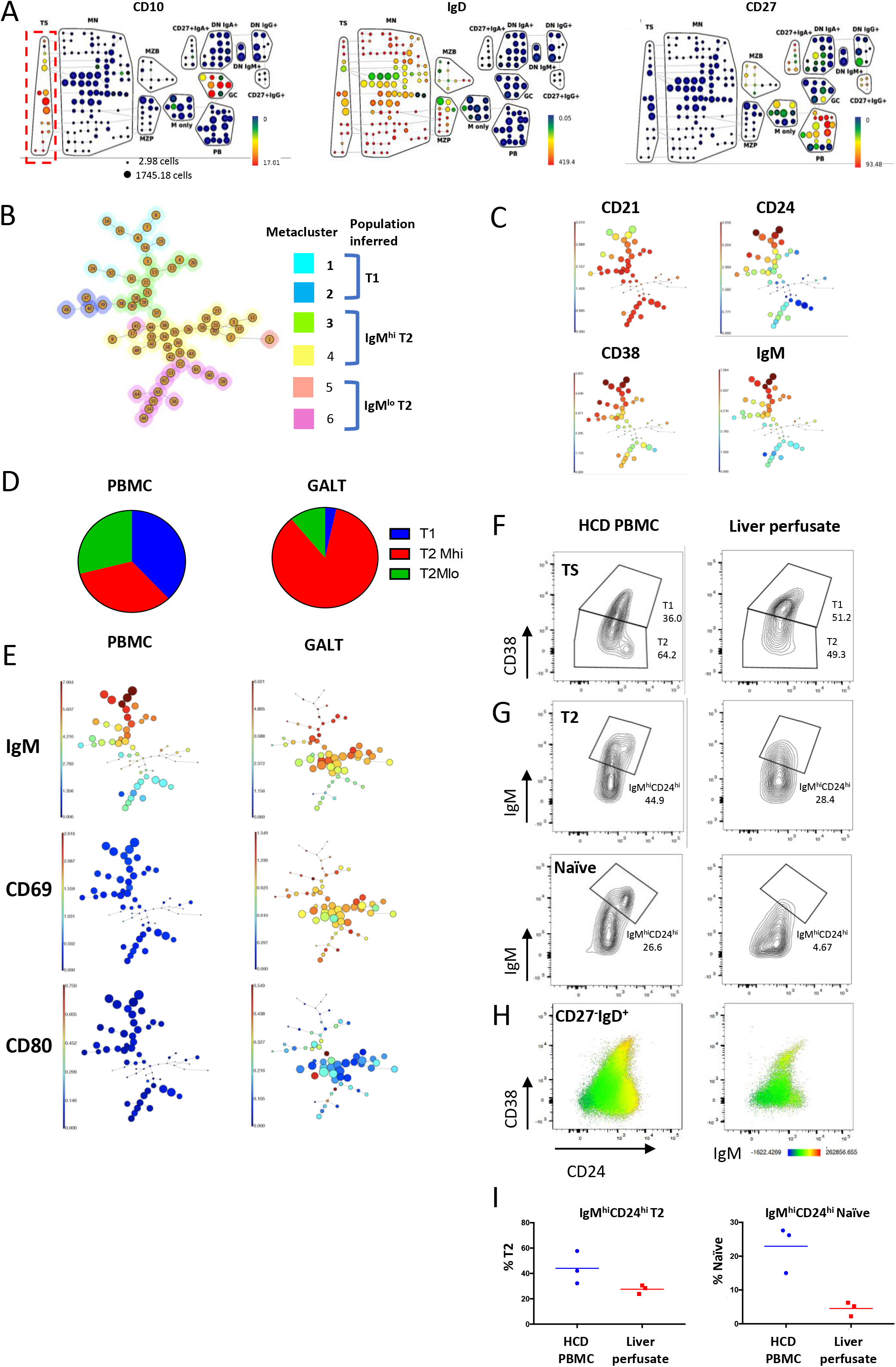
Gut-associated lymphoid tissue is enriched in IgM^hi^ T2 cells. A) SPADE on viSNE plots depicting the expression of B cell lineage markers used to identify T1 and T2 cells as CD27^-^IgD^+^CD10^+^ in a concatenated GALT sample, M only = IgM only memory (CD27^+^IgD^-^IgM^+^), GC = germinal centre (IgD^-^CD10^+^), PB = plasmablast (IgD^-^ CD38^hi^) (see also Fig. S1 H). B) A minimal spanning tree generated by FlowSOM run on exported events (n= 4520) from the TS bubble in Figure part A using CD10, CD24, CD38 and IgM as clustering parameters. Automatic metaclustering of the flowSOM nodes identified 6 metaclusters, the identity of each can be inferred by the relative expression of CD21, CD24, CD38 and IgM (see also Figure part C). C) Minimal spanning trees showing expression of CD21, CD24, CD38 and IgM on a concatenated PBMC sample. D) Pie charts demonstrating the proportion of TS cell subsets inferred from metaclusters in figure part B confirm that GALT is enriched in IgM^hi^ T2 cells. E) Minimal spanning trees demonstrating higher expression of CD69 and CD80 on GALT TS B cells. F) Flow cytometry contour plots of concatenated (n=3) liver perfusate samples and concatenated HCD PBMC (n=3) demonstrating reduced proportion of CD24^++^CD38^++^ T2 cells in liver perfusates. G) Flow cytometry plots of concatenated liver perfusate and PBMC samples demonstrating reduced frequency of IgM^hi^CD24^hi^ T2 and naïve (CD27^-^IgD^+^CD10^-^) cells in liver perfusate samples compared to HCD PBMC. H) Flow cytometry dot plots with IgM MFI overlay of concatenated PBMC and liver perfusate samples demonstrating reduced frequency of IgM^hi^CD24^hi^ TS and naïve cells in liver perfusate samples. I) Scatter plots of flow cytometry data from individual samples gated as in Figure part G demonstrating reduced frequency of IgM^hi^CD24^hi^ TS and naïve cells (CD27^-^IgD^+^CD10^-^) in liver perfusate samples compared to PBMC (median values).

Having observed that IgM^hi^ T2 cells are enriched in GALT we sought confirmation of selective recruitment by asking whether this population is depleted from blood draining the gut via the hepatic portal vein that we isolated from liver perfusion samples. Flow cytometry demonstrated that liver perfusate samples were enriched in T1 cells as reported previously (Vossenkamper et al., 2013)(Fig. 2 F), and that CD24^hi^IgM^hi^ T2 and IgM^hi^ naïve cells were depleted compared to PBMC from HCD (Fig. 2 G, H and I), consistent with their selective recruitment from blood into GALT.

### Transcriptomic analysis of IgM^hi^ and IgM^lo^ TS B cells demonstrates different upstream regulators of phenotype

Having demonstrated contrasting surface phenotypes and migratory capacity of IgM^hi^ and IgM^lo^ TS B cells, we next sought to identify transcriptomic features differing between them and to gain insight into inducers and regulators of these subsets by single cell RNA sequencing. IgM^hi^ and IgM^lo^ TS B cells from 5 HCD were sorted by FACS, pooled and gene expression libraries were prepared using a 10x Genomics single cell 5’ gene expression workflow (Fig. S2 A, B, C and D). In total, 14499 genes were identified in 4268 cells after quality filtering. The non-linear dimension reduction algorithm UMAP (Becht et al., 2018) was run on differentially expressed genes and demonstrated discreet clustering of IgM^hi^ and IgM^lo^ TS B cells (Fig. 3 A). Selected genes from the top 60 differentially expressed genes are illustrated in Fig. 3 B and 3 C. Transcripts encoding *CD1C* and *MZB1* that are expressed by MZB cells, were amongst the most abundantly expressed genes in IgM^hi^ TS B cells. The lupus risk allele and regulator of toll like receptor 9 (TLR9) responses *PLD4* was the most highly differentially expressed gene in IgM^hi^ TS B cells (Gavin et al., 2018). Undirected clustering of pooled IgM^hi^ or IgM^lo^ TS B cells generated clusters that contained predominantly IgM^hi^ or IgM^lo^ TS B cells (Fig. 3 D and E) that shared enrichment of the genes expressed by these cell subsets (Fig. 3 B and F). High expression of IL4R by IgM^lo^ TS and IgM^lo^ naïve B cells was confirmed by flow cytometry (Fig. S2 E and F). Importantly, *KLF2,* which drives murine follicular B cell development, was upregulated in IgM^lo^ TS B cells (Hart et al., 2011). *CCR7* was upregulated in IgM^lo^ TS B cells supporting the higher surface expression that was evident in the mass cytometry analysis. Higher abundance of transcripts encoding L-selectin by IgM^lo^ TS B cells was also confirmed by qPCR using sorted populations (Fig. S2 G and H).

**Figure 3.**
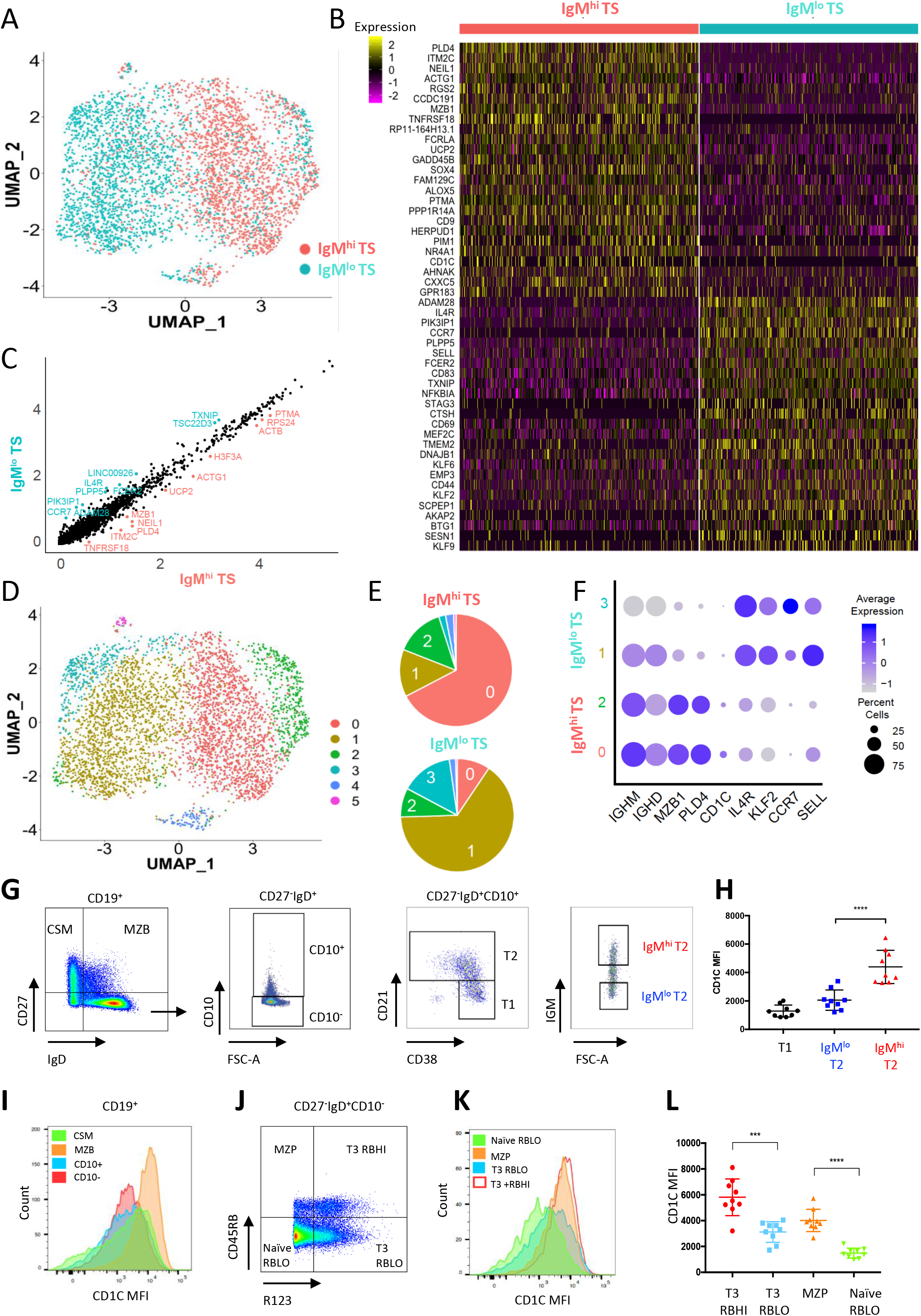
Transcriptomic analysis of IgM^hi^ and IgM^lo^ TS B cells demonstrates different upstream regulators of phenotype. A) UMAP plot of IgM^hi^ and IgM^lo^ TS B cells clustered according to differentially expressed genes identified using the Seurat SCTransform workflow (Hafmeister, 2020) in the two subsets. B) Heatmap of selected genes from the top 60 differentially expressed genes in IgM^hi^ and IgM^lo^ TS B cells. C) Scatter plot demonstrating genes differentially expressed in IgM^hi^ and IgM^lo^ TS B cells. D) A PCA based approach based on differentially expressed genes identified 6 clusters among the IgM^hi^ and IgM^lo^ TS B cells that were demonstrated by UMAP. E) Quantification of the frequency of IgM^hi^ or IgM^lo^ TS B cells within the clusters demonstrated by UMAP in Figure part D reveals that IgM^hi^ TS B cells dominate in clusters 0 and 2 and IgM^lo^ TS B cells in clusters 1 and 3. F) Dot plot demonstrating expression of selected genes within clusters 0-4. G) Flow cytometry dot plots demonstrating the gating strategy to identify IgM^hi^ and IgM^lo^ T2 (CD27^-^IgD^+^CD10^+^CD21^hi^) B cells. H) Scatter plots demonstrating CD1c MFI in T1, IgM^hi^ and IgM^lo^ T2 B cells gated in Figure part G (mean +/− SD, paired t test). I) Histograms demonstrating CD1c MFI in B cell subsets gated in Figure part G. J) Dot plot of flow cytometry data demonstrating the gating strategy used to identify CD45RB^hi^ T3 (R123+) and MZP (R123-) cells. K) A histogram showing CD1c MFI on subsets as gated in Figure part J. L) Dot plots demonstrating CD1c MFI on subsets gated in Figure part J (mean +/− SD, paired t test).

Ingenuity pathway analysis demonstrated enrichment of retinoic acid receptor and lipopolysaccharide induced genes in IgM^hi^ TS B cells (Fig. S2 I and J). IgM^lo^ TS B cells were enriched in genes induced by interferon-γ (IFN-γ), interleukin-1 and interleukin-2 (Fig. S2 K). IgM^hi^ TS B cells used less V_H_1 and more V_H_3 than IgM^lo^ TS B cells consistent with published profiles of MZB repertoire (Bagnara et al., 2015)(Fig. S2 L).

IgM^hi^ and IgM^lo^ TS B cells therefore have distinct transcriptomes. IgM^lo^ cells are selectively enriched in genes encoding peripheral circulation and inhibition of marginal zone B cell fate whereas IgM^hi^ cells have gene expression signatures and IGHV gene family usage linking them to MZB cells.

The abundance of *CD1C* transcripts in IgM^hi^ TS B cells was of particular interest because CD1c is characteristically highly expressed by human MZB (Weller et al., 2004). Consistent with the transcriptomic profile, CD1c surface expression was higher on IgM^hi^ than IgM^lo^ TS B cells (Fig. 3 G and H). As previously reported CD1c expression was high on MZB (Fig. 3 I) as well as on MZP and CD45RB^hi^ T3 cells (previously referred to as T3’) that have been linked to MZB development (Bemark et al., 2013; Descatoire et al., 2014; Koethe et al., 2011; Zhao et al., 2018). MZP and CD45RB^hi^ T3 cells were defined by the phenotype CD27^-^IgD^-^CD10^-^ CD45RB^hi^ with expression of the ABCB1 cotransporter or not, respectively (Fig. 3 J, K and L). Cells that express the ABCB1 cotransporter extrude rhodamine 123 (R123) and are therefore identified as R123^lo^ cells in this analysis. Both subsets share high expression of IgM and CD24 (Fig. S2 M and N)

### Lineage progression from IgM^hi^ TS B cells through to MZB

The shared surface properties of IgM^hi^ TS with IgM^hi^ naïve B cells (Fig. 1 H), the enrichment of transcripts considered characteristic of MZB in IgM^hi^ TS (Fig. 3 B and F), and shared high expression of CD1c by IgM^hi^ TS with MZB and other B cell subsets associated with MZB development (Fig. 3 G-L) all support the existence of an IgM^hi^ MZB differentiation pathway that begins during TS B cell development. We investigated this further by performing pseudotime trajectory analysis of single cell RNA sequencing data from HCD B cells from blood.

CD19^+^ B cells were sorted from PBMC of 3 HCD (Fig. S3 A) and surface labelled with Total-Seq-C antibodies prior to capture on the 10x chromium controller (Fig. S3 B). Gene expression, and antibody detection tag (ADT) libraries were then prepared according to the manufacturer’s instructions and sequenced on an Illumina HiSeq High Output platform (Fig. S3 C).

Data from single HCD were initially analysed individually. UMAP plots were used to visualize clusters and identify the B cell subsets they corresponded to by overlaying signal from lineage defining transcripts and CITE-Seq antibodies (Fig. S3 and S4). TS B cells were identified as CD27^-^IgD^+^ clusters with high surface expression of CD38. Of the remaining CD27^-^IgD^+^ clusters that represented naïve cells, those with the top 30% of median IgM-ADT signal were designated IgM^hi^ (Fig. S3 and S4). Note that because identification of MZP and CD45RB^hi^ T3 would require reagents that are incompatible with this method (Fig. 3 J), they will be included in the IgM^hi^ naïve cell groups in this analysis. CD27^+^IgD^+^ clusters that were enriched in *CD1C* transcripts were designated as MZB. CD27^+^IgD^-^IgM^+^ clusters were designated ‘IgM-only’ cells and CD27^+^IgD^-^IgM^-^ clusters enriched in *HOPX* and *COCH* transcripts were designated as class switched memory B cells (CSM) (Descatoire et al., 2014)(Fig. S3 and S4).

Three dimensional UMAP plots were then used to better visualise the spatial relationship between these B cell subsets (Fig. 4 A, B, C and D and S4 C and F). This demonstrated clear separation of CD27^+^ and CD27^-^ ‘islands’ of cells (Fig. 4 A, B and S4 and Supplemental videos 1, 2 and 3). In all three HCD two distinct cellular ‘bridges’ linked the CD27^-^ and CD27^+^ islands in the plot (Fig. 4 A, B and S4 C and F and Supplemental videos 1, 2 and 3). In each HCD, an IgM^hi^ bridge that was enriched in cells with *CD1C* transcripts linked the CD27^-^ island to MZB. (Fig. 4 A, B, C and D and S4 B and E). In contrast, IgM-only cells were connected to the CD27^-^ island by naïve cells with lower expression of IgM.

**Figure 4.**
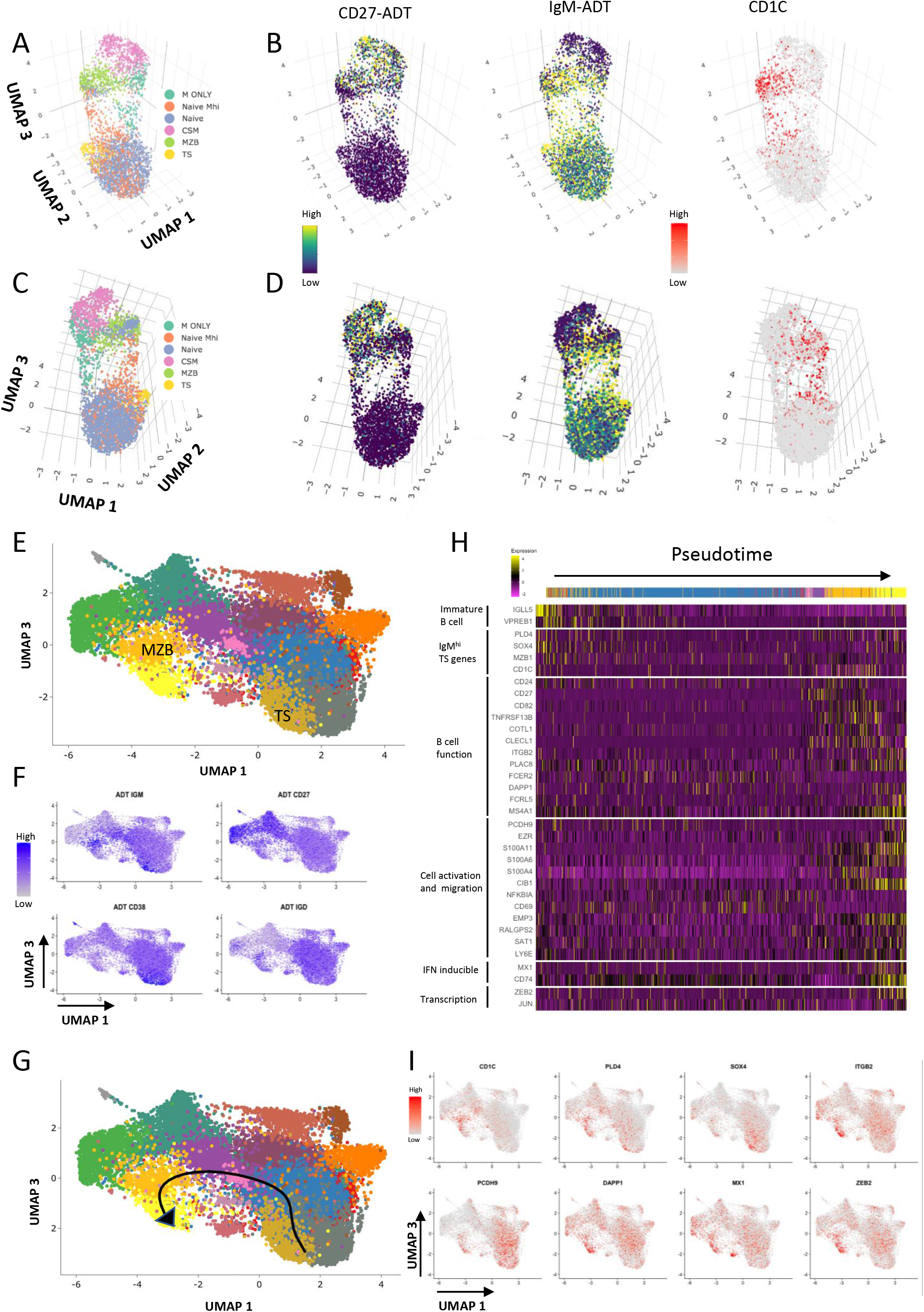
Lineage progression from IgM^hi^ TS B cells through to MZB. A. A 3D UMAP plot of CD19^+^ cells from a HCD (10x HCD1) generated from a PCA run on 2000 differentially expressed genes. Clusters were merged and pseudocoloured according to the B cell subsets they represent as described in Fig. S3 D, E, F and G. B. CD27 and IgM ADT and *CD1C* gene signal overlay on the 3D UMAP plot displayed in Figure part A. C. The 3D UMAP plot depicted in Figure part A viewed using different UMAP axis coordinates to Figure part A. D. CD27 and IgM ADT and *CD1C* gene signal overlay on the 3D UMAP projection in Figure part C. E. A UMAP plot generated by integrating 3 HCD 10x datasets and generated from a PCA run on 2000 variably expressed genes. F. Feature plots demonstrating overlay of ADT signal on the UMAP plot from Figure part E enables identification of clusters representing TS and MZB. G. A Slingshot developmental trajectory overlaid onto the UMAP plot from Figure part E demonstrating developmental progression from clusters representing TS B cells to MZB via IgM^hi^ naïve cells. H. A heatmap of selected genes from the top 100 most differentially expressed genes along the Slingshot trajectory demonstrated in Figure part G. I. Feature plots demonstrating overlay of gene signal from the differentially expressed genes along the Slingshot trajectory identified in Figure part G.

Having visualized the juxtaposition of IgM^hi^ naïve cells with MZB in UMAP clusters we next used the Slingshot tool for pseudotime trajectory analysis. Data from the three HCD were normalized and integrated. UMAP plots were used to identify clusters representing 27^-^ CD38^hi^CD24^hi^ TS cell and CD27^+^IgD^+^IgM^+^ MZB subsets by overlay of CD27, IgM, IgD and CD38 ADT signal (Fig. 4E and F). The TS B cell cluster was selected as the starting point for analysis of pseudotime transitions in Slingshot. Importantly, end points were not specified.

Slingshot identified an IgM^hi^ pseudotime trajectory from TS that passed through the MZB cluster via IgM^hi^ naïve B cells (Street et al., 2018) (Fig. 4 G and Supplemental video 4). Amongst the 100 most differentially expressed genes along this trajectory were *PLD4, CD1C, SOX4* and *MZB1,* that were previously identified as differentially expressed between IgM^hi^ and IgM^lo^ TS B cells (Fig. 3B and F and Fig. 4 H and I). Analysis of gene expression by cells along the trajectory demonstrated progressive downregulation of *IGLL5* and *VPREB1* markers of B cell immaturity (Fig. 4 H). Upregulated in the terminal stages of the trajectory were genes encoding proteins implicated in cell adhesion, including *ITGB2, PCDH9* and activation including *DAPP1* (Fig. 4 H and I). The final cluster in the pseudotime trajectory was enriched in interferon regulated genes *MX1* and transcription factor *ZEB2* (Fig. 4 H and I). IFN induced genes as well as *DAPP1* and *FCRL5* are highly expressed by DN2 cells, although the relationship of this subset with MZB is not known (Jenks et al., 2018). Pseudotime analysis of HCD PBMC therefore identified an IgM^hi^ developmental trajectory from TS B cells to MZB.

### IgM^hi^ and IgM^lo^ TS B cells differ functionally, and in their potential to differentiate

We next determined if IgM^hi^ and IgM^lo^ TS B cells that have different cell surface and transcriptomic characteristics maintain their relative levels of IgM expression *in vitro* following stimulation and if they differ functionally. Initially, proliferation in response to anti-IgM in the presence of CD40L was measured. IgM^hi^ TS B cells proliferated more than IgM^lo^ cells in response to anti-IgM (Fig. 5 A and B). Next we investigated the response of IgM^hi^ and IgM^lo^ TS B cells to the TLR9 agonist CpG, that has been proposed to drive MZB differentiation (Guerrier et al., 2012). In culture, CpG increased surface expression of IgM on both IgM^hi^ and IgM^lo^ B cells. However, IgM^hi^ cells remained IgM^hi^ compared to the IgM^lo^ cells (Fig. 5 C and D). Furthermore, culture with CpG resulted in greater upregulation of CD45RB on IgM^hi^ TS and IgM^hi^ naïve cells than IgM^lo^ TS and IgM^lo^ naïve cells, consistent with adoption of an MZP like phenotype (Fig. 5 E).

**Figure 5.**
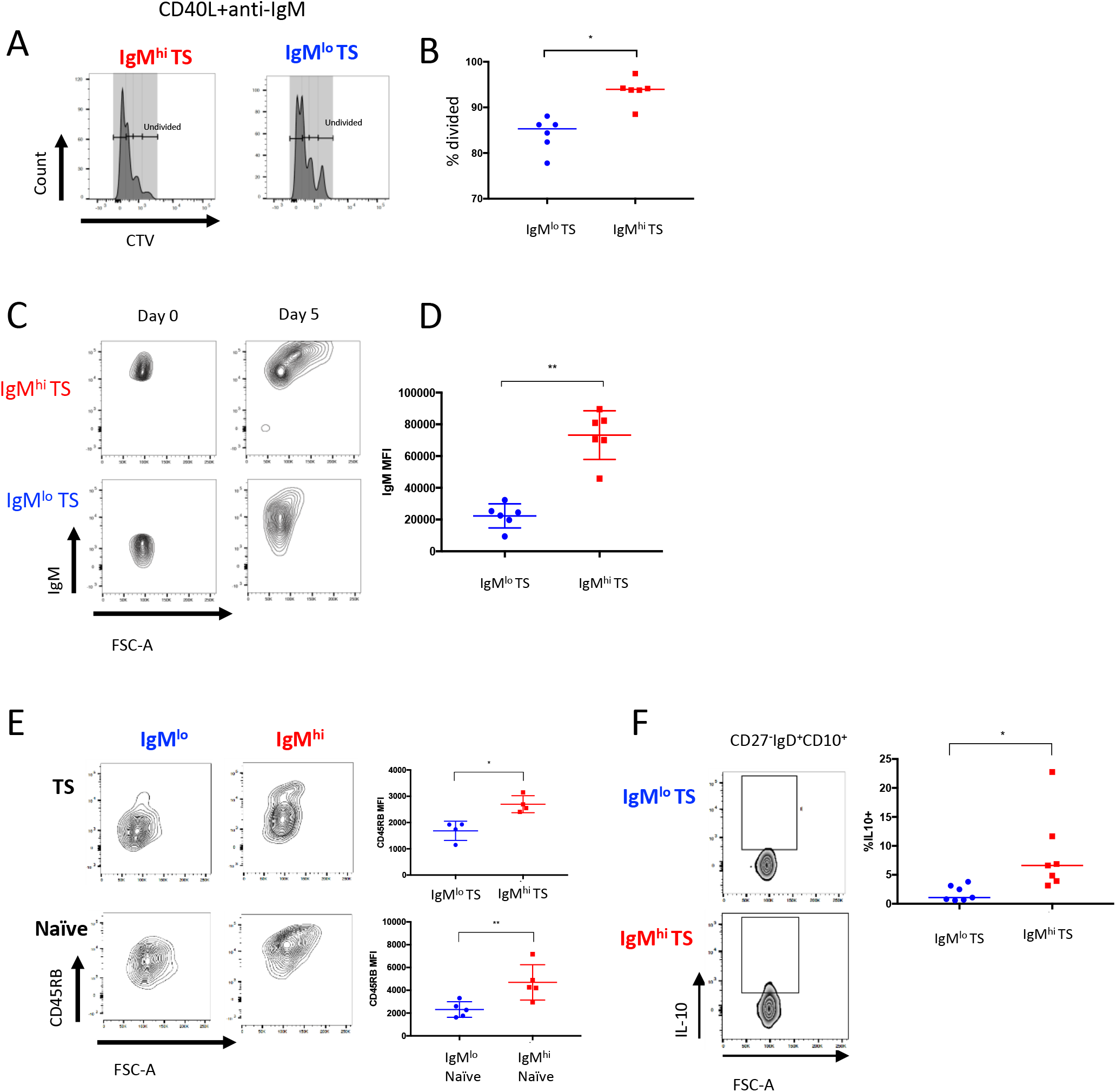
IgM^hi^ and IgM^lo^ TS B cells differ functionally and in their potential to differentiate. A) Histograms of flow cytometry data demonstrating cell trace violet (CTV) dilution following 5 days of culture with CD40L+anti-IgM in IgM^hi^ and IgM^lo^ TS B cells. B) Scatter plots demonstrating increased proliferation of IgM^hi^ TS compared to IgM^lo^ TS B cells when stimulated with CD40L+anti-IgM for 5 days (medians, Wilcoxon test). C) Flow cytometry contour plots demonstrating that an IgM^hi^ phenotype is maintained on IgM^hi^ TS B cells after CpG stimulation for 5 days. D) Scatter plots demonstrating that an IgM^hi^ phenotype is maintained on IgM^hi^ TS B cells after CpG stimulation for 5 days (mean +/− SD, paired t test). E) Flow cytometry contour plots and scatter plots demonstrating greater upregulation of CD45RB by IgM^hi^ than IgM^lo^ TS and naïve (CD27^-^IgD^+^CD10^-^) subsets at day 5 culture with CpG (mean +/− SD, paired t test). F) Flow cytometry contour plots and scatter plots demonstrating higher frequency of IL10 expressing cells among IgM^hi^ TS B cells following 6 hour stimulation with PMA/ionomycin (median, Wilcoxon test).

A subpopulation of human cells with a TS phenotype are regulatory, and murine IL10 producing B regulatory (Breg) cells are T2 marginal zone progenitor cells and the gut is important for their induction (Blair et al., 2010; Pillai et al., 2005; Rosser et al., 2014). We therefore investigated the capacity of IgM^hi^ TS B cells to produce IL10. Following 6 hours stimulation with PMA and ionomycin, IgM^hi^ TS B cells produced significantly more IL10 than IgM^lo^ cells (Fig. 5 F), inferring greater regulatory capacity of this subset.

### Marginal zone B cell differentiation is defective in patients with severe SLE

We have previously observed reduced frequencies of circulating TS B cells expressing β7 integrin in a subset of SLE patients, implying reduced potential for TS B cells to access GALT in these cases. Data presented here implicates GALT as an important site for MZB differentiation and MZB depletion has been reported in SLE (Rodriguez-Bayona et al., 2010; Zhu et al., 2018). Hence, we sought to determine whether our proposed MZB differentiation pathway was defective in SLE.

Flow cytometry was used to quantify B cell subsets in a cohort of 41 SLE patients and matched HCD (Table S1,2). Reduced MZB frequency was seen in patients with SLE compared to HCD (Fig. 6 A and B) and this was most marked in patients with lupus nephritis (LN) compared to patients with other manifestations of SLE (OL) (Fig. 6 C). Reduced MZB frequency was not a feature of other autoimmune diseases studied (Fig. 6 C, Table S3), although it has been identified in patients with Sjögren’s disease (Roberts et al., 2014). We found that a relative reduction of MZB in patients with SLE was associated with a reduction of MZP (CD27^-^IgD^+^CD10^-^CD45RB^hi^ R123^-^) (Fig. 6 D and E), T3 CD45RB^hi^ cells (CD27^-^IgD^+^CD10^-^ CD45RB^hi^R123^+^)(Fig. 6 D and F). This was again most consistently observed in the LN patient cohort. The proportion of naïve B cells (CD27^-^IgD^+^CD10^-^R123^lo^) was also diminished in SLE (Fig. 6 G) but CD45RB^lo^R123^hi^ cells were more frequent (Fig. 6 H). This population was further divided into T3 and aNAV by their expression of CD24 and CD38 (Fig. 6 I). Whilst both subsets were increased in patients with LN, CD45RB^lo^R123^hi^ cells were predominantly T3 cells (Fig. 6 I). Frequencies of MZB correlated with both MZP and T3 CD45RB^hi^ frequencies and inversely correlated with T3 CD45RB^lo^ frequencies (Fig. 6 J).

**Figure 6.**
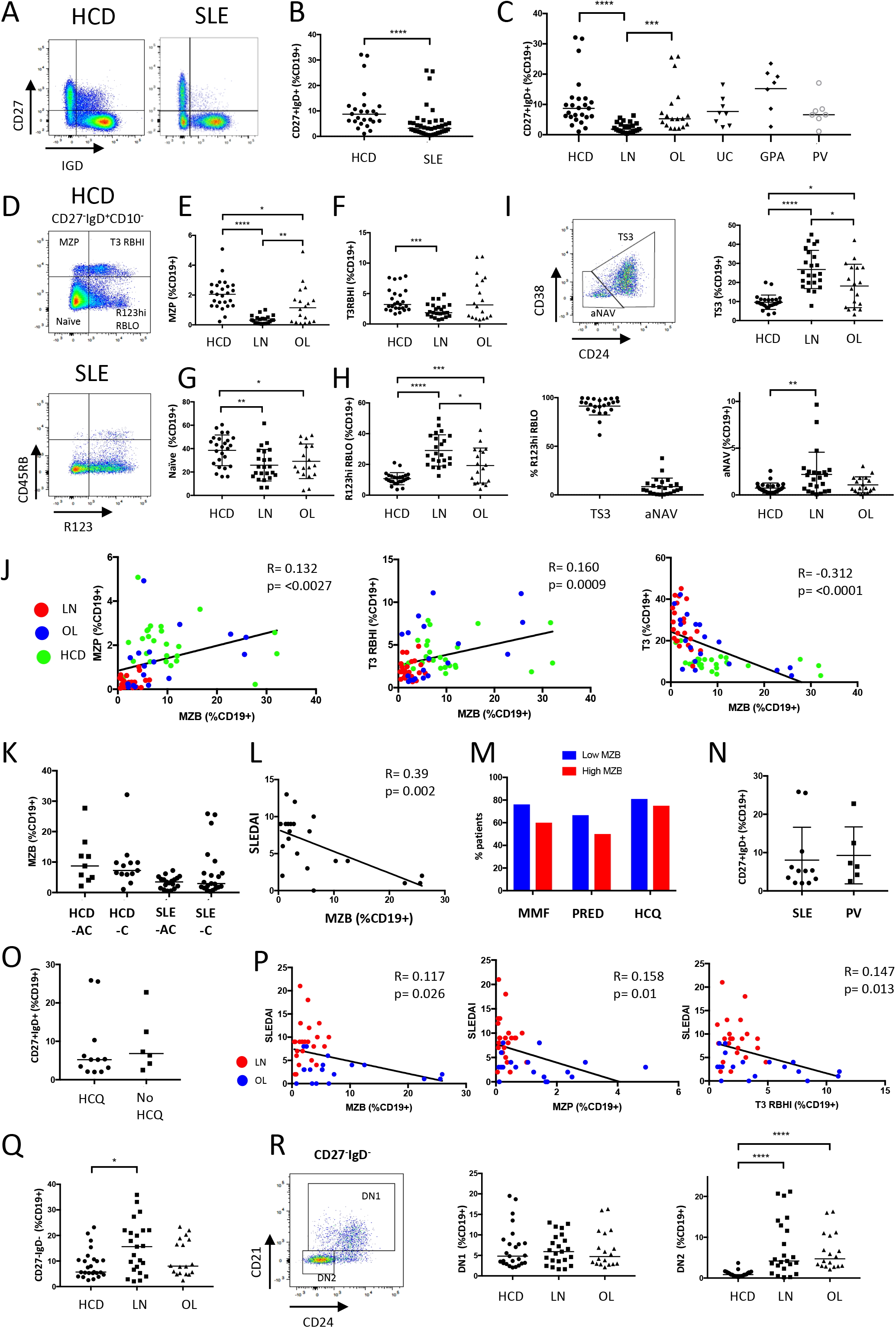
Marginal zone B cell differentiation is defective in patients with severe SLE. A) Flow cytometry dot plots and scatter plots demonstrating reduced frequency of CD27^+^IgD^+^ (MZB) cells in a patient with SLE compared to a HCD. B) Scatter plots of flow cytometry data demonstrating reduced frequency of MZB in patients with SLE compared to HCD (medians, Mann-Whitney test). C) Scatter plots of flow cytometry data demonstrating reduced MZB frequency in lupus nephritis (LN), other lupus subtypes (OL) but not in ulcerative colitis (UC), granulomatosis with polyangiitis (GPA) and pemphigus vulgaris (PV) (medians, Mann-Whitney test). D) Flow cytometry dot plots of a HCD and an SLE patient demonstrating identification of T3 (R123^hi^) and naïve (R123^lo^ subsets with high and low expression of CD45RB. Stark reduced frequency of CD45RB^hi^ T3 and naïve (MZP) populations is evident in SLE. E) Scatter plot of flow cytometry data demonstrating reduced frequency of MZP (CD45RB^hi^R123^lo^) cells in LN patients (medians, Mann-Whitney test). F) Scatter plot of flow cytometry data demonstrating reduced frequency of CD45RB^hi^ T3 (R123^hi^) cells in LN patients (medians, Mann-Whitney test). G) Scatter plot of flow cytometry data demonstrating reduced frequency of naïve (CD45RB^lo^R123^lo^) cells in LN patients (mean +/− SD, unpaired t test). H) Scatter plot of flow cytometry data demonstrating enrichment of CD45RB^lo^R123^hi^ cells in LN patients (mean +/− SD, unpaired t test). I) Gating of CD24^+^CD38^+^ T3 and CD24^-^CD38^-^ aNAV cells revealed that CD45RB^lo^R123^hi^ cells were mostly T3 cells, and that it was this population that was enriched in LN patients. J) The proportion of MZB showed a positive correlation with the proportion of MZP and T3 CD45RB^hi^ cells but a negative correlation with T3 (CD45RB^lo^R123^hi^CD24^+^CD38^+^) cells (Spearman’s rank coefficient). K) Scatter plots of flow cytometry data demonstrating no difference in MZB frequency in African Caribbean (-AC) and Caucasian (-C) HCD and SLE patients (medians, Mann-Whitney test). L) Correlation of MZB and SLEDAI score in Caucasian SLE patients (Spearman’s rank coefficient). M) Bar graphs demonstrating the immunosuppressive burden of SLE patients with low MZB counts (< 3.13% CD19+ cells) vs high MZB counts (>3.13% CD19+ cells) where 3.13% represents the median MZB value in all SLE patients. N) Scatter plots of flow cytometry data demonstrate that pemphigus vulgaris (PV) patients taking MMF and / or prednisolone did not have reduced MZB when compared to SLE patients on the same immunosuppressive medication (mean +/− SD, unpaired t test). O) Scatter plots of flow cytometry data demonstrate there was no difference in MZB counts in non-renal SLE (OL) patients taking or not taking hydroxychloroquine (HCQ) therapy (medians, Mann-Whitney test). P) The proportion of MZB, MZP and T3 CD45RB^hi^ showed a negative correlation with disease activity as indicated by the SLEDAI score (Spearman’s rank coefficient). Q) Scatter plots showing the proportion of CD27^-^IgD^-^ B cells as gated in Figure part A demonstrate increased frequency of this population in LN (medians, Mann-Whitney test). R) Flow cytometry dot plot demonstrating the identification of DN1 and DN2 cells based on expression of CD21 and CD24, and scatter plots showing that DN2 cells were more abundant in LN than in HCD (medians, Mann-Whitney test).

We next asked if reduced MZB frequency in LN patients was intrinsic to the disease or secondary to patient demographics, immunosuppressants or disease severity. HCD, LN and OL cohorts were matched for age, gender and ethnicity (Table S1,2) and there was no difference in MZB frequency between Caucasian and African Caribbean healthy donors and SLE patients (Fig. 6 K). A subset of Caucasian SLE patients were seen to have higher MZB than African-Caribbean patients but this subset had mild disease (Fig. 6 L). There was no difference in immunosupression between SLE patients with high and low MZB frequencies and MZB depletion was not seen in patients with pemphigus vulgaris (PV) taking prednisolone and mycophenolate mofetil (MMF) and did not differ between SLE patients taking or not taking hydroxychloroquine (HCQ) (Fig. 6 M, N and O). However, MZB, MZP and CD45RB^hi^ T3 cell frequencies correlated with disease severity (Fig. 6 P). The more marked reduction of MZB in the LN patients is therefore likely to be due to this patient group representing a severe disease cohort. As previously reported, CD27^-^IgD^-^ DN cells were more abundant in LN (Fig. 6 Q). These were predominantly CD24^lo^CD21^lo^ and therefore DN2 consistent with other studies (Fig. 6 R)(Jenks et al., 2018).

MZB depletion in SLE is therefore associated with reduced frequency of MZP and T3 CD45RB^hi^ cells. This consolidates the concept of these cells as being in a developmental continuum in health and suggests that aberrant transitional B cell maturation may result in failure of their genesis in SLE.

### IgM^hi^ β7 integrin^hi^ T2 cells are reduced in frequency in lupus nephritis

To identify early stages of aberrant marginal zone lineage development in LN we used mass cytometry to compare blood B cell subsets from LN patients and HCD in an undirected way (Fig. S5 A, B and C). The automated clustering algorithm CITRUS identified populations that differed significantly in abundance between LN and HCD. The three main clusters of nodes (Fig. 7 A) can be identified by their relative expression of B cell lineage markers (Fig. 7 B and C). CD27^-^IgD^+^CD10^-^ CD45RB^hi^ (MZP) and CD27^+^IgD^+^CD10^-^ CD45RB^hi^ (MZB) cells were significantly depleted in LN patients (Fig. 7 Ci and Cii). TS B cells were more abundant (Fig. 7 Ciii) whilst IgA class switched cells were depleted in patients with LN (Fig. 7 Civ).

**Figure 7.**
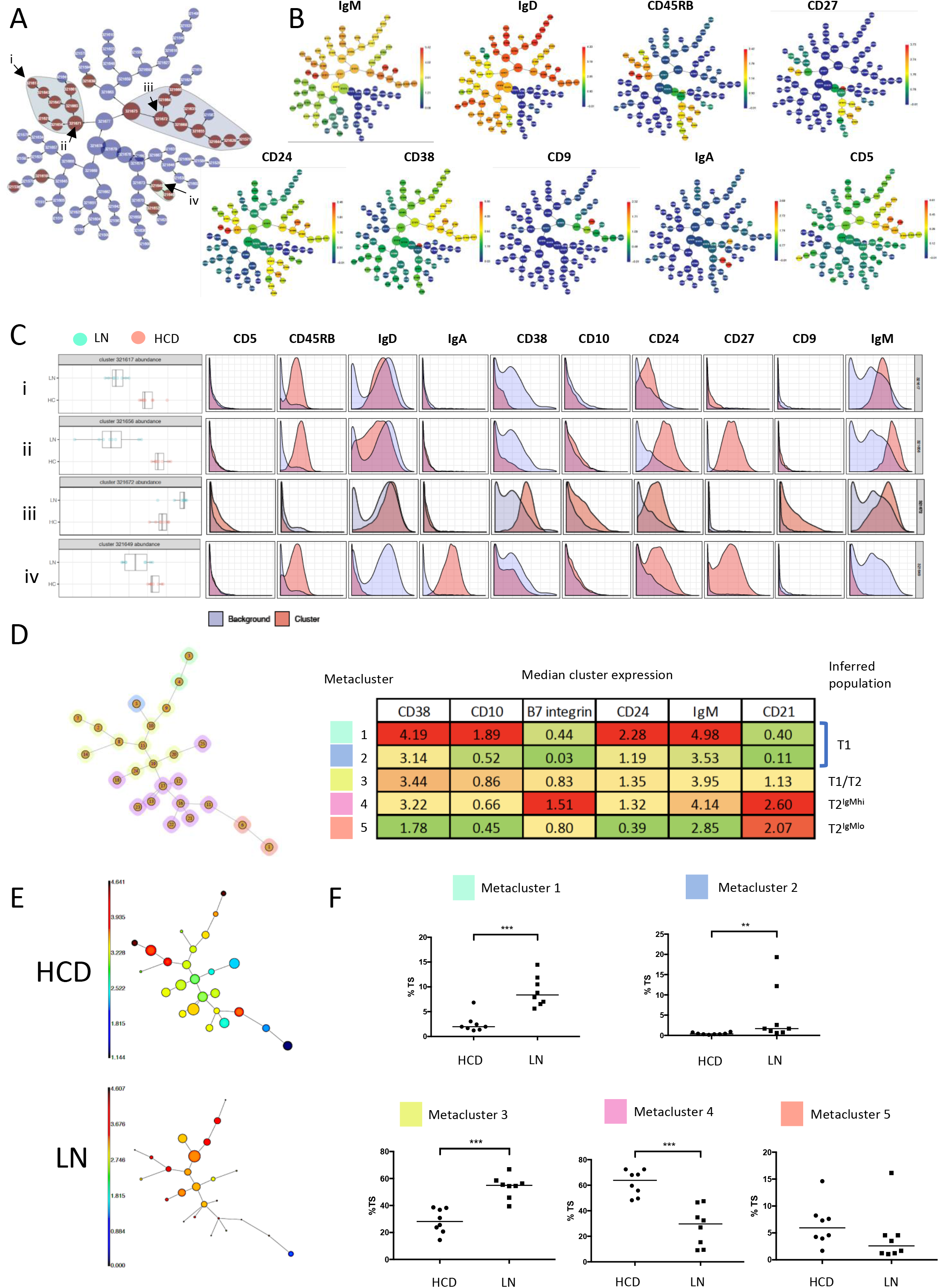
IgM^hi^ β7 integrin^hi^ T2 cells are reduced in frequency in lupus nephritis. A) CITRUS trees generated from CD19+ cells from HCD (n=8) and LN patients (n=8) and clustered according to the expression of CD5, CD9, CD10, CD24, CD27, CD38, CD45RB, IgD, IgM and IgA (see also Fig. S5 A, B and C, Table S1). Red nodes indicate significantly different population abundances between HCD and LN patients. A grey background is automatically assigned to aggregates of significant nodes. Arrows and roman numerals indicate nodes further analysed in Figure part C. B) CITRUS trees demonstrating the median expression of the clustering panel markers in the nodes. C) Histograms demonstrating the abundance and expression of panel markers in selected nodes, the identify of these nodes can be inferred as (i) MZP, (ii) MZB, (iii) TS B cells and (iv) class switched IgA memory. D) Minimal spanning tree generated by FlowSOM from events exported from CITRUS node (iii) identified in Figure parts A and C. This generated 5 metaclusters, the identification of each could be inferred by median expression of panel markers in each metacluster. E) Minimal spanning trees generated by FlowSOM plots demonstrating CD38 expression in a representative HCD and LN patient. Prominent skewing of TS cell subpopulations is evident in LN. F) Relative abundances of metaclusters as a percentage of TS B cells indicate reduced frequency of events within metacluster 4 corresponding to IgM^hi^ β7^hi^ TS B cells in LN, and increased frequency of events in metaclusters 1,2 and 3 corresponding to T1 cells (medians, Mann-Whitney test).

TS B cell subpopulations in HCD and LN were quantified by FlowSOM (Fig. 7 D and E). 5 metaclusters were identified representing subdivisions of T1 and T2 populations (Fig. 7 D). Inspection of minimal spanning trees demonstrated stark depletion of certain TS subpopulations in LN (Fig. 7 E). Quantification of events within metaclusters from all donors revealed that T1 cells represented by metaclusters 1,2 and 3 were more abundant in LN whilst IgM^hi^ T2 cells with high expression of β7 integrin represented by metacluster 4 were markedly less abundant in LN patients (Fig. 7 F).

Depletion of IgM^hi^ T2 cells with high expression of β7 integrin is therefore associated with defective MZB maturation in LN patients. This affirms the association between MZB and IgM^hi^ T2 cells in health and implicates reduced access of these cells to GALT in the breakdown of this developmental axis in patients with severe lupus (Fig. S5 D).

## Discussion

We have identified branches of human B cell lineage maturation that are evident from the T2 stage. An IgM^hi^ branch, that expresses higher levels of β7 integrin and lower levels of IL4R compared to the IgM^lo^ branch, is gut homing. Confirmation of differentiation through IgM^hi^ stages of differentiation from IgM^hi^ T2, including IgM^hi^CD45RB^hi^ T3 and naïve B cell variants to MZB is gained from pseudotime analysis coupled with the observed concerted reduction of the stages in this sequence in patients with severe SLE (Fig. S5 D). The reduced frequencies of gut homing IgM^hi^ T2 cells in severe SLE further consolidates the role of GALT in early B cell fate decisions and supporting MZB development. The collapse of this gut homing MZB maturational axis in severe SLE therefore affirms its existence in health as well as having important disease relevant implications.

We have previously observed that human T2 cells are recruited into GALT where they are activated by intestinal microbes (Vossenkamper et al., 2013). Here we demonstrate that specifically the IgM^hi^ T2 subset of TS B cells is recruited into GALT, where they have a phenotype of activated cells including expression of CD69 and CD80. The IgM^hi^ T2 subset is also enriched in ROR and LPS inducible genes, consistent with exposure to the microbiota. We show that the TLR9 agonist CpG that upregulates IgM and NOTCH2 in human TS B cells (Capolunghi et al., 2008; Guerrier et al., 2012) also upregulates CD45RB on IgM^hi^ TS and IgM^hi^ naïve cells. *PDL4,* that is a lupus risk allele and the most highly upregulated gene in IgM^hi^ compared to IgM^lo^ TS B cells, is upregulated along the developmental pathway to MZB and limits responses to CpG (Gavin et al., 2018). This suggests that PDL4 defects could contribute to SLE pathogenesis by impacting an aspect of the development or function of IgM^hi^ TS B cells involving TLR9. Interestingly, *PLD4* is also expressed in the splenic marginal zone in mice (Yoshikawa et al., 2010) and *PLD4* knockout mice develop autoantibodies and immune complex mediated renal damage similar to SLE with LN (Gavin et al., 2018). IgM^hi^ TS B cells also show a transcriptomic signature indicative of retinoic acid regulation that is a feature of GALT microenvironment. Together, these data suggest that innate signals and the gut environment impact the origin, fate and function of IgM^hi^ TS B cells.

Consistent with proposed developmental continuum from the IgM^hi^ T2 stage through to MZB, GALT is involved in MZB development, including a stage of receptor diversification in GALT GC. However, supporting a relatively short-term transit coupled to differentiation, the frequencies of somatic mutations in MZB are lower than those of memory B cells or plasma cells in the gut (Zhao et al., 2018). Together these data suggest that GALT transit and GC occupancy are important but transient phases in IgM^hi^ T2 to MZB lineage progression. Pseudotime analysis also identified a population of B cells that appear to develop from MZB and that are activated and more mature. It is possible that activation of MZB might generate a novel population of effector or memory cells.

MZB differentiation is associated with a distinctive gene expression changes and acquisition of the transcription factor ZEB2 (SIP1). ZEB2 has previously been identified as a component of a network including miR200 and TGF-β1 that can regulate cell fate decisions (Gregory et al., 2008; Guan et al., 2018). Activated TGFβ-1 is produced abundantly in the gut. It is possible that in addition to playing important roles in regulation of intestinal immunity as a switch factor for IgA and induction of regulatory T cells, it could also be involved in gut-associated MZB development by interactions with ZEB2 (Borsutzky et al., 2004; Chen et al., 2003).

Collapse of the MZB developmental pathway in severe SLE was accompanied by expanded T3, aNAV and DN2 cell populations. Expansion of aNAV and DN2 populations is a product of excessive TLR7 and IFN-γ signalling. We were therefore interested in enrichment of IFN-γ induced genes in IgM^lo^ TS B cells. Interestingly, the IFN-γ regulated transcription factor KLF2 was transcriptionally upregulated in IgM^lo^ TS B cells. KLF2 drives follicular B cell maturation in mice and its deletion results in an expansion of MZB cells. The role of KLF2 in human B cell development is not known, however loss-of-function *KLF2* mutations along with *NOTCH2* mutations that increase the stability of the notch intracellular domain (NICD) are the most commonly encountered mutations in human MZB cell lymphoma (Campos-Martin et al., 2017). This implicates KLF2 in human B cell fate decisions and eludes to a role for IFN-γ in B cell development and supports its proposed involvement of the imbalance of B cell subsets in LN. The role of IFN-γ in defective MZB maturation is also supported by depletion of this subset in patients with severe COVID19, which is associated with elevated serum IFN-γ levels and extrafollicular B cell repsonses (Laing et al., 2020; Woodruff et al., 2020).

LN represents a severe lupus subtype associated with the worst clinical outcomes (Yap et al., 2012). B regulatory (Breg) IL10 responses associated with expression of CD80 and CD86 are defective in SLE (Blair et al., 2010), permitting aberrant T effector functions (Oleinika et al., 2019). In mice Breg cells are IgM^hi^CD21^hi^CD23^hi^ T2 MZP cells and interaction with the gut microbiome is essential for their induction (Evans et al., 2007; Rosser et al., 2014). We have identified that IgM^hi^ TS B cells express CD80 in GALT and represent the predominant IL10 producing TS B cell subset. Their depletion in LN may be synonymous with the loss of Breg IL10 responses and associated with the lack of T cell regulation in SLE. MZB confer immunity to encapsulated bacteria such as pneumococcus, thus their depletion in LN may confer increased risk of such infections SLE (Danza and Ruiz-Irastorza, 2013). This also reinforces the importance of pneumococcal vaccination in this patient cohort.

In summary, we identify an MZB maturation pathway that becomes evident at the T2 stage of B cell development and that is depleted in severe SLE. Traffic through GALT is a component of this pathway that is potentially linked to the induction of human IL10 producing Breg cells (Rosser et al., 2014). Together, this affirms the importance of tissue microenvironments in shaping the B cell functional repertoire and maintaining health. Understanding the regulators of early B cell fate will be a key to resolving the disturbances in B cell function in severe SLE.

## Acknowledgements

We thank sample donors clinical research support staff L.Nel and N.Morton. This work was funded by the Medical Research Council of Great Britain (MR/R000964/1, MR/L009382/1, MR/P021964/1 and MR/R000964/1) and The St Thomas’ Lupus Trust. We acknowledge financial support from the Department of Health via the National Institute for Health Research comprehensive Biomedical Research Centre award to Guy’s & St Thomas’ NHS Foundation Trust and King’s College London for the flow and mass cytometry at the Flow Cytometry Research Platform, and library preparation and sequencing at the Genomics Research Platform. MB is supported by research funds from The Swedish Research Council and the County Council of Västra Götaland.

## Author contributions

Conceptualization and design of study: T.J.T., Y.Z., D.D’C., M.B., J.S.; Sample identification and collection: T.J.T, W.G., Y.Z., W.J., M.D.R, R.W.G., J.D.S., D.D’C.; Data acquisition and methodology; T.J.T., C.L-F., W.G., Y.Z., U.D.K., P.D., S.J., R.E., S.H., M.B., J.S.; Data analysis: T.J.T, M.J.P, W.G., J.H.S, N.P., S.H, R.E, M.B.; Supervision and funding: J.S., M.B., D.D’C.

## Materials and Methods

### Lead contact

Information regarding reagents and resources should be directed to Professor Jo Spencer.

### Data and code availability

The datasets generated during this study will be made available on acceptance of the manuscript.

## EXPERIMENTAL SUBJECT DETAILS

All blood and tissue samples were obtained from adults with REC approval and informed consent. SLE patients were recruited using the following criteria; i) fulfilment of 4 or more revised ACR classification criteria; ii) ANA positive; iii) Biologic (Belimumab or rituximab) naïve; iv) Immunosuppressive regimen does not include azathioprine or cyclophosphamide within 6 months of sample collection due to the severe depletion of naïve B cells by these medications. All LN patients had diagnostic confirmation by renal biopsy. Blood was obtained from SLE patients and HCD (REC reference 11/LO/1433: Immune regulation in autoimmune rheumatic disease). Paired gut biopsies and blood were obtained from individuals undergoing colonoscopies in whom no mucosal abnormality was detected (REC reference 11/LO/1274: Immunology the intestine; features associated with autoimmunity). Samples of draining the gut via the hepatic portal vein were obtained from liver perfusion prior to transplantation (REC reference 09/H0802/100: The role of innate immune system in hepatic allograft outcome). Patient demographic data can be found in Tables S1 and S2.

## METHODS

### Sample Processing

Blood samples were diluted 1:1 in RPMI-1640 containing 10% foetal calf serum (FCS), 100 U/ml penicillin and 100 μg/ml streptomycin (RPMI-P/S). Diluted blood was then layered onto Ficoll and centrifuged for 25 minutes with brake and accelerator set to 0. The buffy coat layer was then removed and cells were washed in RPMI-P/S. PBMC isolated from patients undergoing colonoscopy was used fresh, whilst PBMC used for the analysis of HCD and patients with SLE, UC, GPA and PV was cryopreserved in FCS + 10% dimethyl sulfoxide (DMSO). Mononuclear cells from gut were obtained by the removal of epithelial cells with 1mM EDTA in HBSS containing 100 U/ml penicillin and 100 μg/ml streptomycin for 30 minutes. Collagenase digest was then used to generate a cell suspension using collagenase D (1 mg/ml) and DNase (10 U/ml) in RPMI-P/S for 1 h.

### Mass cytometry

3 mass cytometry panels were utilized, the staining protocols were as follows.

Panel 1: Cryopreserved cells were washed and rested in RPMI-P/S + 0.1mg/ml DNase at 37 degrees for 45 minutes. B cells were then negatively enriched using a Miltenyi B cell isolation kit II. 4 x 10^6^ were then viability stained with 1ml cisplatin 25μM in 1x PBS. Cells were then washed in PBS containing 0.5% BSA with 2mM EDTA (Cell staining medium, C-SM) and resuspended in 10ul Fc receptor blocking solution and left for 10 minutes on ice. IgG staining was then performed in 100μl staining volume for 30 minutes on ice. Cells were then washed in CS-M and resuspended in the pre-titrated volume of antibody mastermix, the volume was then adjusted to 100ul with CS-M and cells were stained for 30 minutes on ice. Metal tagged antibodies used are listed in Fig S1 and The Key Resources Table. Cells were then washed twice in 1xPBS and fixed overnight in 16% paraformaldehyde. The following day cells were washed in 1xPBS and DNA was stained with 1μM Intercalatin in 500ul permeabilization buffer at room temperature for 20 minutes. Cells were then washed twice in 1xPBS and twice in Milli-Q water before being resuspended in Milli-Q water plus EQ beads to a concentration of 0.5 x 10^6^ /ml and run on a Helios Mass cytometer.

Panel 2: As for panel 1 except cells were stained fresh, were not enriched and 2 x 10^6^ cells were viability stained with 1ml rhodium intercalator diluted in 1:500 in PBS for 20 minutes at room temperature. Metal tagged antibodies used are listed in Fig S2 and The Key Resources Table.

Panel 3: As per panel 1 except cells were not enriched and IgG staining was not performed. Metal tagged antibodies used are listed in Fig S6 and The Key Resources Table.

### Analysis of mass cytometry data

FCS files were normalised using Nolan lab software (v0.3, available online at https://github.com/nolanlab/beadnormalization/releases). Pre and post normalisation plots are shown in Fig. S1 B and F and S5 A for the respective datasets. Where files were concatenated the Cytobank FCS File Concatenation Tool was used (available online at https://support.cytobank.org/hc/en-us/articles/206336147-FCS-file-concatenation-tool). Files were then loaded onto the Cytobank (https://mrc.cytobank.org/) and gated to identify live CD19 B cells and analysed as described in Fig. S1 C and G and S5 C.

For the analysis of HCD PBMC in Fig. 1, viSNE was run on equal numbers of CD19^+^ events (n=35000) from each HCD (n=10). SPADE was then run on the viSNE coordinates and B cell subsets were identified by placing nodes into bubbles. The TS bubble was identified as CD27^-^IgD^+^CD24^+++/++^CD38^+++/++^. Events within the TS bubble were exported and a further viSNE was run using equal events (n=3535) and all panel markers except CD45, CD3, CD14 and class switched isotypes IgA and IgG which are not expressed by TS B cells. CD45 was excluded due to homogenous expression and lack of contribution to clustering. SPADE was then run on the viSNE coordinates and TS populations were defined as demonstrated in Fig. 1 B.

For the analysis of PBMC and GALT derived B cells in Fig. 2, equal numbers of CD19^+^ events (n=118,934) from concatenated PBMC (n=7) and GALT (n=7) samples were used to run a viSNE using all markers except for CD45, CD3 and CD14. SPADE was then run on the viSNE coordinates and TS B cells identified as CD27^-^IgD^+^CD10^+^ nodes. Events within the TS bubble were then exported and equal numbers of events (n=4520) were clustered using FlowSOM. CD10, CD24, CD38, IgM as clustering channels to allow the undirected visualization of markers of TS populations.

For the analysis of PBMC from HCD and SLE samples in Fig 7, CITRUS was run using equal numbers of CD19^+^ events (n=20000) from HCD (n=8) and SLE patients (n=8) and the following clustering channels : CD5, CD9 CD10, CD24, CD27, CD38, CD45RB, IgD, IgM, IgA. Due to event sharing amongst CITRUS nodes, node 321672 identified in Fig. 7 A contains all CD27^-^IgD^+^CD24^+++/++^CD38^+++/++^ events and was therefore used for analysis of TS B cells. All events from this node were exported and FlowSOM was run using equal event sampling (n= 657) and using all marker channels except CD45, CD3, CD14 and IgA.

### Flow cytometry and cell sorting

Cryopreserved cells used for flow cytometry were thawed and washed in RPMI-P/S and then rested at 37 degrees in RPMI-P/S + 0.1mg/ml DNase for 45 minutes. Viability staining with Zombie aqua dye was performed using 100μl 1:200 dilution in 1xPBS, or with DAPI 0.1mg/ml diluted 1:1000 and added prior to sample acquisition on the flow cytometer. Cells were stained on ice for 15 mins with pre-titrated concentrations of antibodies listed in The Key Resources Table. Staining with R123 was performed for 10 minutes at a concentration of 6μM and cells were washed and chased for 3 hours in RPMI-P/S. All samples were analysed by a BD LSRFortessa (BD Biosciences). Anti-Mouse/Rat beads (BD) were used for fluorescent compensation and gates were set using appropriate isotype controls. Cell sorting was performed using a BD FACSAria (BD Biosciences) and live single CD19^+^ B cells were gated as follows IgM^hi^ TS : CD27^-^IgD^+^CD10^+^IgM^hi^, IgM^lo^ TS : CD27^-^IgD^+^CD10^+^IgM^lo^, IgM^hi^ naïve: CD27^-^IgD^+^CD10^-^IgM^hi^, IgM^lo^ naïve : CD27^-^IgD^+^CD10^-^IgM^lo^, where IgM^hi^ and IgM^lo^ gates captured 30% of the highest and lowest IgM expressing cells respectively.

### Cytokine detection

Fresh PBMC was isolated from HCD and incubated for 6 hours at 37 degrees with phorbol 12-myristate 13-Acetate (PMA) 50ng/ml and Ionomycin 250ng/ml with Golgiplug at a dilution of 1:1000. Cells were then surface stained as above followed by fixation with Cytofast buffer (Biolegend). Cells were then washed twice and stained with resuspended conjugated antibodies in permeabilization / wash buffer (Biolegend) for 20 minutes at room temperature.

### Cell culture and stimulation analysis

Sorted IgM^hi^ and IgM^lo^ TS and naïve (CD27^-^IgD^+^CD10^-^) B cell subsets were plated onto a 96 well plate seeded with 2 x 10^4^ cells per well. Wells containing CD40L expressing HEK cells were also seeded with 2 x 10^4^ irradiated HEK cells per well. Cells were then stimulated with CPG-ODN 2.5μg/ml or anti-IgM 10μg/ml. Proliferation assays were performed on cells stained with CellTrace violet as per the manufacturer’s guidelines. Cells were then stained and analysed by flow cytometry as above.

### Single cell RNA sequencing library preparation

Sorted cell populations were loaded onto a 10x Genomics Chromium Controller and 5’ gene expression, VDJ and ADT (for samples in Fig. 4) were prepared according to the manufacturers guidelines. Samples used in Fig. 3 were sequenced using an Illumina NextSeq 500 platform. Samples used in Fig. 4 were sequenced using an Illumina HiSeq 2500 High Output platform. The 10x Genomics Cellranger workflow was then used for transcript alignment and the generation of sparse matrices for downstream analysis.

### CITE-Seq antibody staining

Cryopreserved samples were thawed and sorted using the gating strategy in Fig. S3 A. Cells were then washed and stained in a CITE-Seq antibody cocktail at a concentration of 8μg/ml for 30 minutes on ice. Cells were then washed three times before loading onto the 10x Chromium controller.

### Single cell sequencing analysis

The Seurat R package (vs 3.1.1) was used to filter data to remove cells with low numbers of RNA transcripts, doublets and cells with high levels of mitochondrial transcripts indicative of cell death. Immunoglobulin variable genes were then removed from the dataset as well as cells with low expression of B cell genes *CD79A, CD79B, CD19* or *MS4A1.* Data from IgM^hi^ and IgM^lo^ TS B cells were merged and the data was transformed in accordance with the SCTransform workflow before UMAP based reduction of dimensionality and PCA-based clustering to identify populations (Hafmeister, 2020). Heatmaps were then created using select genes from the top 60 differentially expressed genes in each sample, and dot plots and violin plots on selected genes. Data from sorted CD19^+^ cells from HCD used for Fig. 4 and Fig. S3 and 4 was initially analysed individually followed by an integrated analysis. Individual analysis was performed using the quality control (QC) steps as well as the removal of IGHV genes and non-B cells as described above. Data was then normalized and scaled and UMAP run on a PCA generated using 2000 variable genes. Overlay of ADT and gene signal, violin plots and median expression of markers by UMAP clusters was used to identify which B cell subsets they corresponded to. For the integrated data analysis, data from 3 HCD was filtered using the QC steps as well as the removal of IGHV genes and non-B cells as described above. Data was then normalized using the SCTransform wrapper in Seurat followed by integration using The Satija Laboratory Integration and Label Transfer protocol (Butler et al., 2018), using 3000 integration features. The 2000 most variable genes were then used to perform PCA and a 3-dimensional UMAP was obtained from this.

Clusters were obtained using the FindNeighbours and FindClusters functions within Seurat, using default parameters. The UMAP coordinates and cluster allocations were then used to run Slingshot (Street et al., 2018). Randomised downsampling of 50% was required to improve the performance of trajectory inference in Slingshot. ADT overlay of the UMAP plot was used to identify the cluster composed of CD27-IgD+CD38^hi^ cells that best represented TS B cells and this was chosen as the starting point from which Slingshot would build trajectories. A heatmap was then created using genes of interest amongst the top 100 differentially expressed genes on the trajectory.

### Quantitative RT-PCR

Quantitative RT-PCR was performed using Taqman Gene Expression Assays (FAM, Thermofisher Scientific) were used to quantify CCR7 and ITGB7 expression in cDNA from sorted IgM^hi^ and IgM^lo^ TS B cell subsets. Reactions were performed in duplicate and multiplexed with Eukaryotic 18S rRNA Endogenous Control (VIC). Samples were run on a QuantStudio 5 Real Time PCR System (Thermofisher Scientific). △CT was calculated using Thermofisher Connect software (available online at (https://apps.thermofisher.com/apps/spa/#/dataconnect).

## QUANTIFICATION AND STATISTICAL ANALYSIS

### Flow cytometry and mass cytometry data

Flow cytometry data was visualized and gated using FlowJo v 10.6.1. Mass cytometry data was analysed using cytobank software.

### Statistical analysis

Graphpad prism version 7.0 was used for statistical analysis. Paired t tests or Wilcoxon tests were used to compare paired samples whilst unpaired t tests or Mann-Whitney tests were used for unpaired samples. Adjusted p values were represented as * p=<0.05, ** p=<0.01, *** p=<0.001. All error bars show the mean +/− standard deviation.

## Supplemental videos

Supplemental video 1. Rotation of the 3D UMAP plot demonstrating B cell subsets from 10x

HCD1 as depicted in Fig. 4A and S3 D, E, F and G.

Supplemental video 2. Rotation of the 3D UMAP plot demonstrating B cell subsets from 10x

HCD2 as demonstrated in Fig. S4 A, B and C.

Supplemental video 3. Rotation of the 3D UMAP plot demonstrating B cell subsets from 10x

HCD3 as demonstrated in Fig. S4 D, E and F.

Supplemental video 4. Rotation of 3D UMAP plot as depicted in Fig. 4 G with overlay of IgM ADT signal demonstrating that the Slingshot trajectory passes through IgM^hi^ naïve B cells.

---

**Fig. S1.**
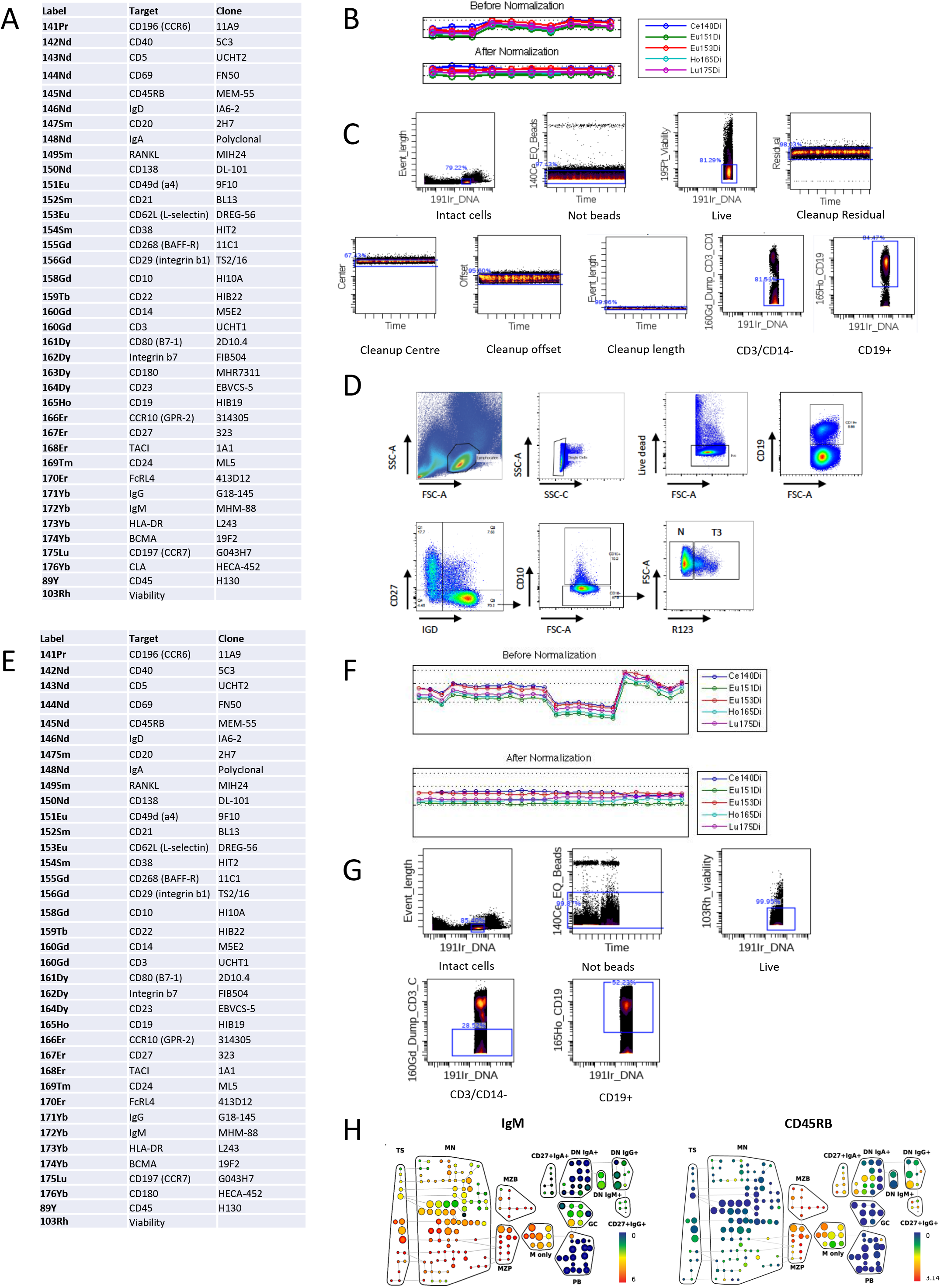
Gating and analysis of mass cytometry data used for Fig. 1 and 2. **A)** Mass cytometry panel used for analysis in Figure 1. **B)** Pre- and post normalization plots of mass cytometry data used for Fig.1. **C)** Gating strategy of mass cytometry data to identify live CD19^+^ B cells. **D)** Flow cytometry plots demonstrating identification of T3 as CD27^-^IgD^+^CD10^-^R123^hi^ and naïve (N) B cells as CD27^-^ IgD^+^CD10^-^R123^lo^. **E)** Mass cytometry panel used for analysis in Figure 2. **F)** Pre- and post normalization plots of mass cytometry data used for Fig. 2. **G)** Gating strategy of mass cytometry data to identify live CD19^+^ B cells. **H)** Spade trees demonstrating expression of IgM and CD45RB in the concatenated GALT sample (see also Fig. 2 A.)

**Fig. S2.**
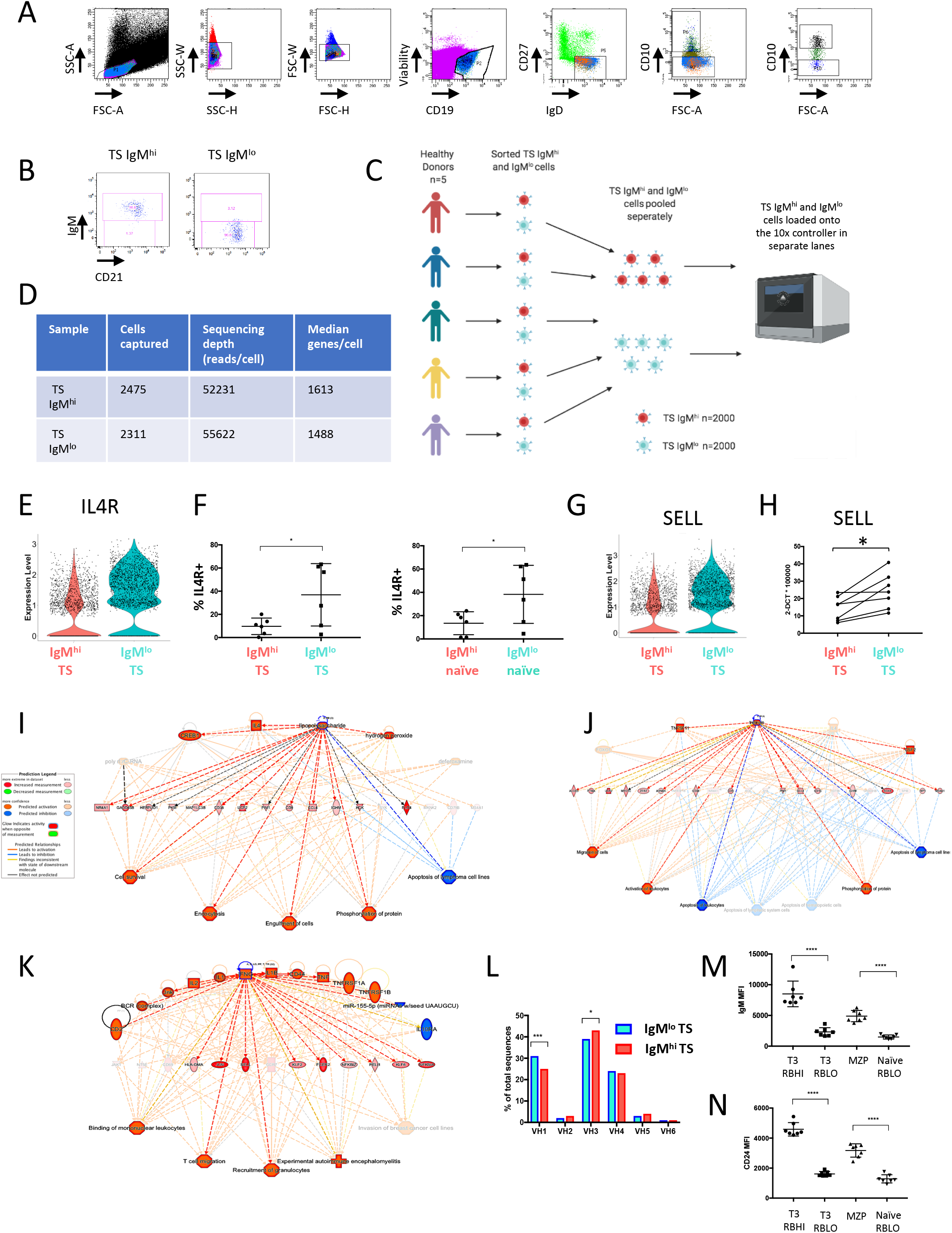
Sort strategy, 10x genomics workflow and validation. **A)** FACS sort strategy to identify IgM^hi^ and IgM^lo^ TS B cell subsets. **B)** Purity plots of sorted IgM^hi^ and IgM^lo^ TS B cell subsets. **C)** 10x genomics experimental workflow detailing pooling of HCD samples. **D)** Summary table of cell numbers captured by the 10X controller and sequencing depth of IgM^hi^ and IgM^lo^ TS B cell subsets. **E)** Violin plot demonstrating expression of the *IL4R* gene in IgM^lo^ TS B cells. **F)** Scatter plots of flow cytometry data demonstrating higher frequency of IL4R on IgM^lo^ compared to IgM^hi^ TS (CD27^-^IgD^+^CD10^+^) and naïve (CD27^-^IgD^+^CD10^-^) cells (mean +/− SD, paired t test). **G)** Violin plot demonstrating expression of the *SELL (CD62L)* gene in IgM^lo^ TS B cells. **H)** qPCR confirms higher levels of the *SELL* gene transcript in IgM^hi^ TS B cells expressed as ΔCT values relative to an 18S endogenous control (paired t test). **I)** IPA upstream regulator plot demonstrating enrichment of LPS induced genes in IgM^hi^ TS B cells. **J)** IPA upstream regulator plot demonstrating enrichment of retinoic acid induced genes in IgM^hi^ TS B cells. **K)** IPA upstream regulator plot demonstrating enrichment of IFN-” induced genes in IgM^lo^ TS B cells. **L)** Bar graphs demonstrating a lower frequency of V_H_1 and higher frequency of V_H_3 immunoglobulin variable heavy chain gene usage in IgM^hi^ TS B cells than TS IgM^lo^ cells (Chi squared test with Bonferroni correction). **M)** Scatter plot of flow cytometry data from HCD demonstrating that T3 and naïve CD45RB^hi^ subsets as gated in Figure 3J share high IgM expression (MFI mean +/− SD, paired t test). **N)** T3 and naïve CD45RB^hi^ subsets share similar high surface expression of CD24 (MFI mean +/−SD, paired t test).

**Fig. S3.**
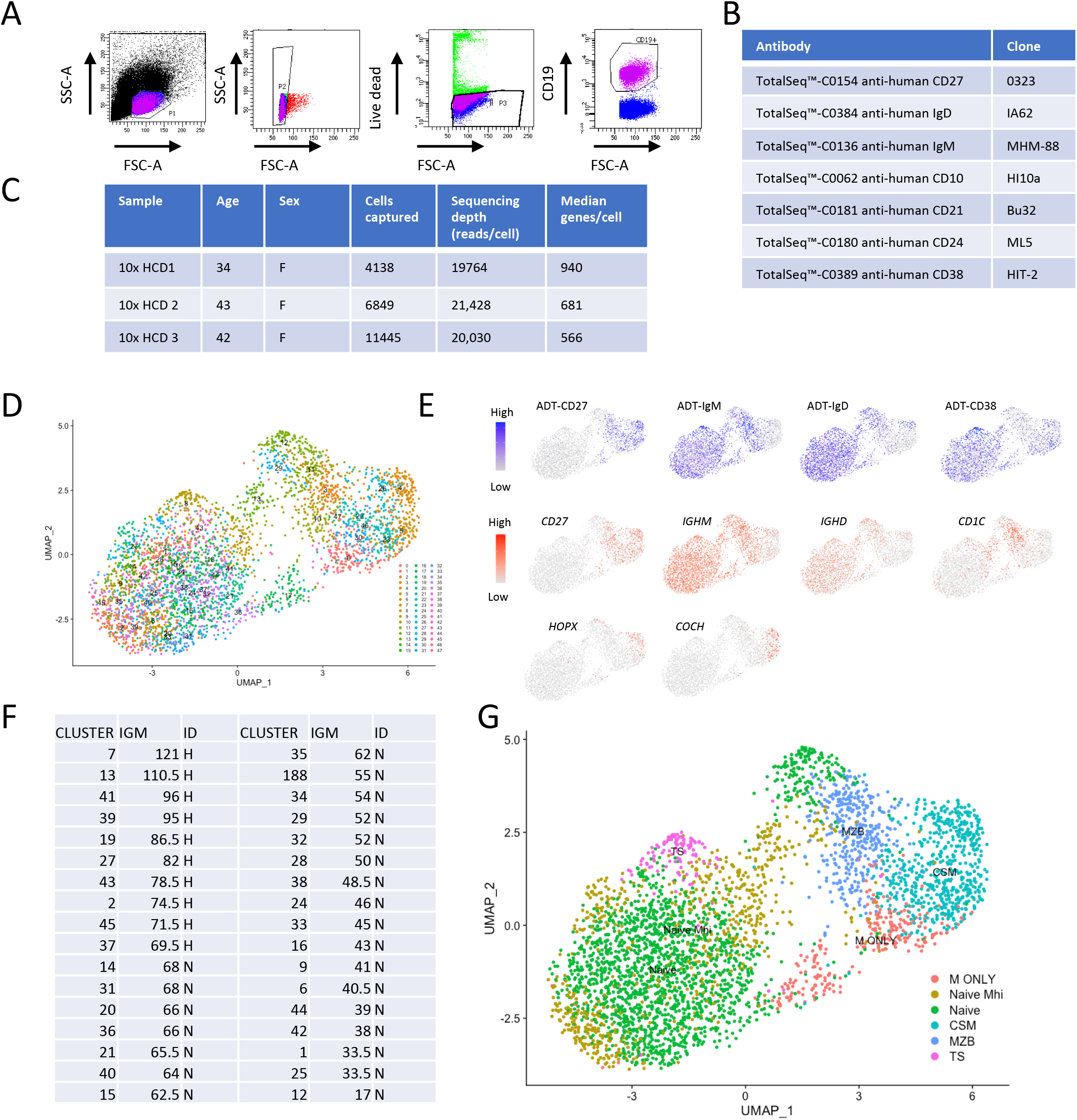
Sort strategy and 10x genomics workflow and identification of B cell subsets represented by UMAP clusters in 10x HCD 1. **A)** Gating strategy to sort live CD19^+^ cells. **B)** Total-Seq antibodies and clones used for surface labelling of CD19^+^ B cells. **C)** Demographic details of HCD, cells captured and sequencing depth. **D)** UMAP plot demonstrating clusters generated from a PCA run on 2000 differentially expressed genes from 10x HCD1. **E)** Feature plots demonstrating lineage defining ADT (CITE-Seq antibody) and transcript signal overlay on the UMAP plot. **F)** Table of median IgM expression within clusters representing naïve cells (CD27^-^IgD^+^CD38^int^), the top 30% of clusters were designated as IgM^hi^. **G)** Merged and pseudocoloured clusters representing B cell subsets defined by ADT and gene signal of lineage defining targets.

**Fig. S4.**
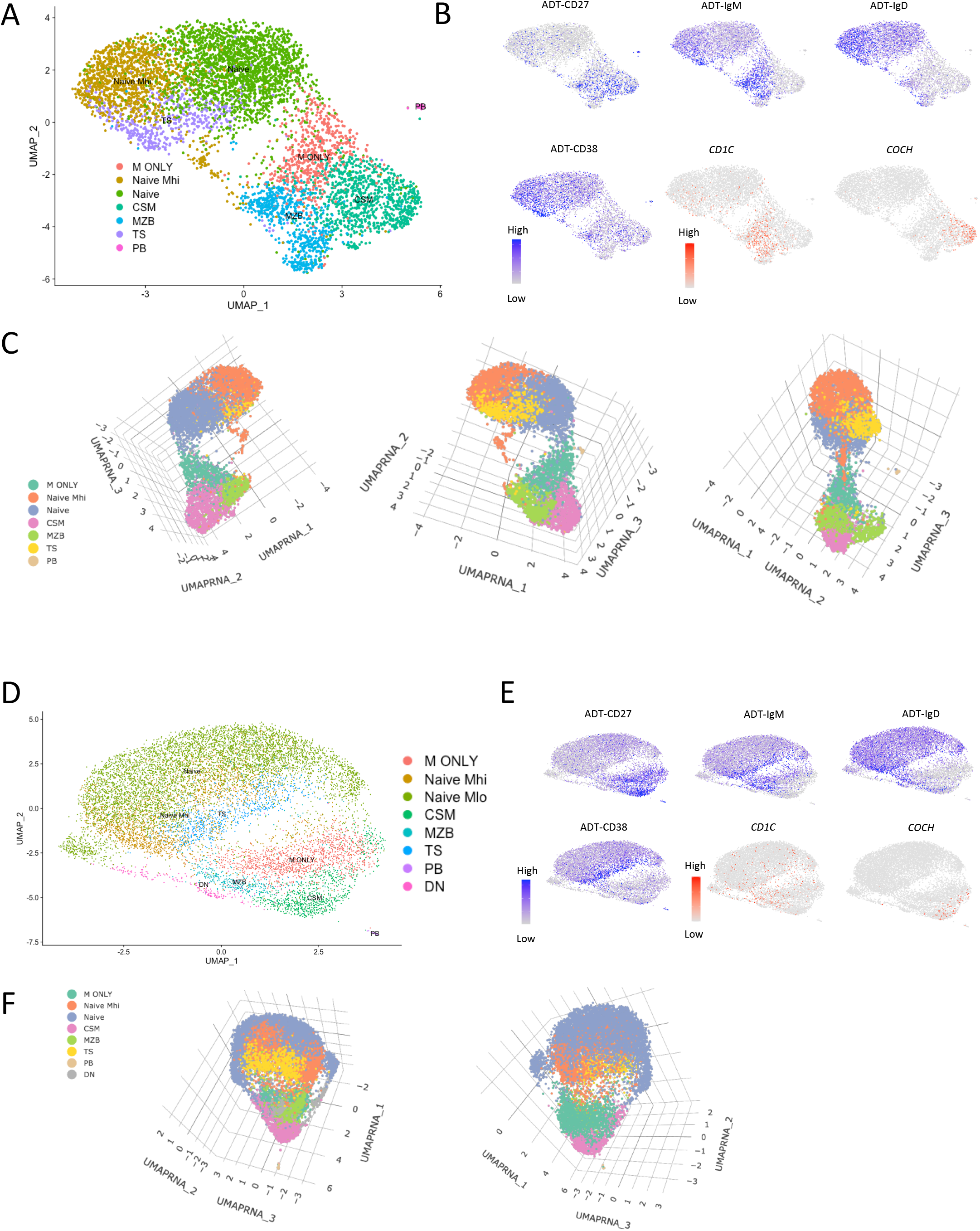
Identification of B cell subsets represented by UMAP clusters in 10x HCD 2 and 3. **A)** UMAP plot generated from a PCA run on 2000 differentially expressed genes from 10x HCD 2. Clusters were merged and pseudocoloured to represent B cell subsets defined by ADT and gene signal of lineage defining targets as described in Figure S4D-G. **B)** Feature plots demonstrating lineage defining ADT (CITE-Seq antibody) and transcript signal overlay on the UMAP plot in Figure part A. **C)** 3D UMAP plots demonstrating the spatial relationship of clusters identified in Figure part A. **D)** UMAP plot generated from a PCA run on 2000 differentially expressed genes from 10x HCD 3. Clusters were merged and pseudocoloured to represent B cell subsets defined by ADT and gene signal of lineage defining targets as described in Figure S4D-G. **E)** Feature plots demonstrating lineage defining ADT (CITE-Seq antibody) and transcript signal overlay on the UMAP plot in Figure part D. **F)** 3D UMAP plots demonstrating the spatial relationship of clusters identified in Figure part D.

**Fig. S5.**
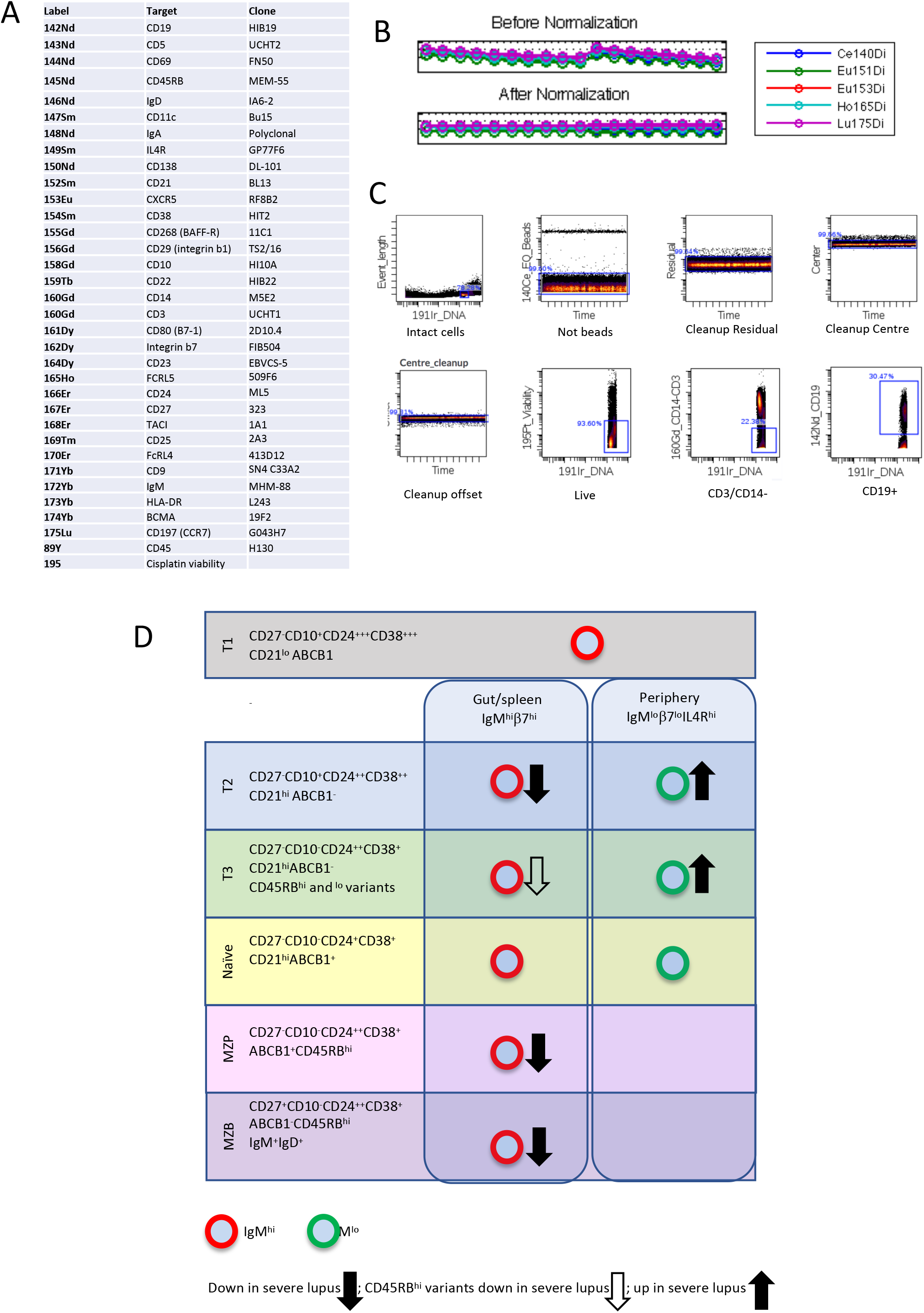
Gating and analysis of mass cytometry data used for Fig. 7 and proposed model of defective MZB differentiation in severe SLE. **A)** Mass cytometry panel used for the analysis in Fig.7. **B)** Pre- and post normalization plots of mass cytometry data used in Fig. 7. **C)** Gating strategy to identify live CD19^+^ B cells. **D)** Proposed model of defective MZB differentiation in severe SLE.

**Table S1.** SLE patient demographic data used for Figure 6 and 7. C = Caucasian, AC = African Caribbean, SEA = South East Asian, MMF = mycophenolate mofetil, PRED = prednisolone, HCQ = hydroxychloroquine.

| Age | Sex | Ethnicity | Nephritis | Cutaneous | Arthritis | Antibodies | Medication | SLEDAI | Figure |
| --- | --- | --- | --- | --- | --- | --- | --- | --- | --- |
| 50 | F | C | + | - | - | ANA, dsDNA | Pred, MMF, HCQ | 9 | 6,7 |
| 63 | f | AC | + | - | - | ANA, dsDNA, Ro, RNP, Sm | MMF, HCQ | 2 | 6,7 |
| 49 | F | AC | + | + | + | dsDNA, Ro | PRED, MMF, HCQ | 9 | 6 |
| 51 | F | C | + | + | + | ANA, dsDNA, RNP | PRED, MMF, HCQ | 2 | 6 |
| 24 | F | C | + | - | - | dsDNA, Sm, RNP | PRED, MMF, HCQ | 6 | 6,7 |
| 30 | F | AC | + | - | - | ANA, dsDNA, la, RNP, Sm | PRED, MMF, HCQ | 7 | 6 |
| 46 | F | AC | + | - | - | ANA, RNP, Sm | MMF, HCQ | 4 | 6 |
| 31 | F | C | + | - | - | ANA, dsDNA | Pred, MMF, HCQ | 9 | 6 |
| 31 | F | AC | + | + | + | ANA, C1Q, dsDNA, RNP, Sm | None | 21 | 6,7 |
| 48 | F | C | + | + | + | ANA, dsDNA, RNP | HCQ | 13 | 6 |
| 45 | M | C | + | - | - | ANA, Sm | PRED, MMF, HCQ | 9 | 6 |
| 30 | f | C | + | - | - | ANA, dsDNA | Pred, HCQ | 7 | 6,7 |
| 27 | F | C | + | + | - | ANA, dsDNA, C1Q | Pred, MMF, HCQ | 9 | 5 |
| 43 | F | AC | + | + | + | ANA, dsDNA, RNP, Sm | Pred, MMF, HCQ | 18 | 6 |
| 53 | F | C | + | + | - | ANA, C1Q, dsDNA, RNP, Ro | PRED, MMF | 9 | 6 |
| 41 | F | C | + | - | - | ANA, dsDNA | PRED, MMF, HCQ | 12 | 6,7 |
| 31 | F | C | + | - | - | ANA, dsDNA | PRED, MMF, HCQ | 5 | 6 |
| 36 | F | AC | + | + | + | ANA, dsDNA, Ro | PRED, MMF, HCQ | 8 | 6 |
| 29 | F | AC | + | - | - | ANA, C1Q, dsDNA, La, RNP, Ro, Sm | MMF, HCQ | 4 | 6 |
| 25 | F | AC | + | - | - | ANA, dsDNA, Ro | PRED, MMF, HCQ | 10 | 6 |
| 68 | F | AC | + | - | - | ANA, dsDNA, RNP, Sm | PRED, HCQ | 13 | 6 |
| 48 | F | C | + | - | - | ANA, dsDNA, RNP, Sm | PRED, MMF, HCQ | 8 | 6 |
| 20 | F | C | + | + | - | ANA, dsDNA, Sm | PRED, MMF, HCQ | 10 | 6 |
| 57 | F | AC | + | - | - | ANA, Sm, RNP | MMF, HCQ | 5 | 7 |
| 48 | F | C | + | - | - | ANA, dsDNA | MMF, PRED, HCQ | 13 | 7 |
| 36 | F | C | - | + | - | ANA, RNP, Ro, Sm | PRED |  | 6 |
| 27 | M | SEA | - | + | - | dsDNA, RNP, Sm | PRED, MMF, HCQ | 4 | 6 |
| 30 | F | AC | - | + | + | Sm, RNP | HCQ | 10 | 6 |
| 59 | f | AC | - | - | + | dsDNA | PRED. | 5 | 6 |
| 35 | F | AC | - | + | + | ANA, dsDNA, La, Ro, Sm | PRED, HCQ | 13 | 6 |
| 56 | F | AC | - | + | - | ANA, RNP, Ro | HCQ | 8 | 6 |
| 53 | F | AC | - | + | - | ANA, RNP, Sm | NONE | 3 | 6 |
| 54 | M | AC | - | - | + | ANA, dsDNA, RNP, Sm | HCQ | 3 | 6 |
| 43 | F | C | - | + | - | ANA, C1Q, dsDNA | PRED, HCQ | 8 | 6 |
| 22 | F | AC | - | + | - | ANA, RNP | PRED, MMF, HCQ | 3 | 6 |
| 52 | F | AC | - | + | - | ANA, dsDNA, Ro | PRED, MMF, HCQ | 0 | 6 |
| 52 | F | C | - | + | + | ANA | NONE | 0 | 6 |
| 58 | F | AC | - | - | + | ANA, Ro | MEP | 3 | 6 |
| 29 | F | C | - | - | + | ANA, dsDNA | PRED, HCQ | 3 | 6 |
| 53 | F | C | - | - | - | ANA, Ro | PRED | 4 | 6 |
| 21 | F | C | - | + | + | ANA, Ro | PRED, MMF | 6 | 6 |
| 30 | F | C | - | - | + | ANA | HCQ | 0 | 6 |
| 39 | F | C | - | - | - | ANA, dsDNA | MMF, HCQ | 2 | 6 |

**Table S2.**
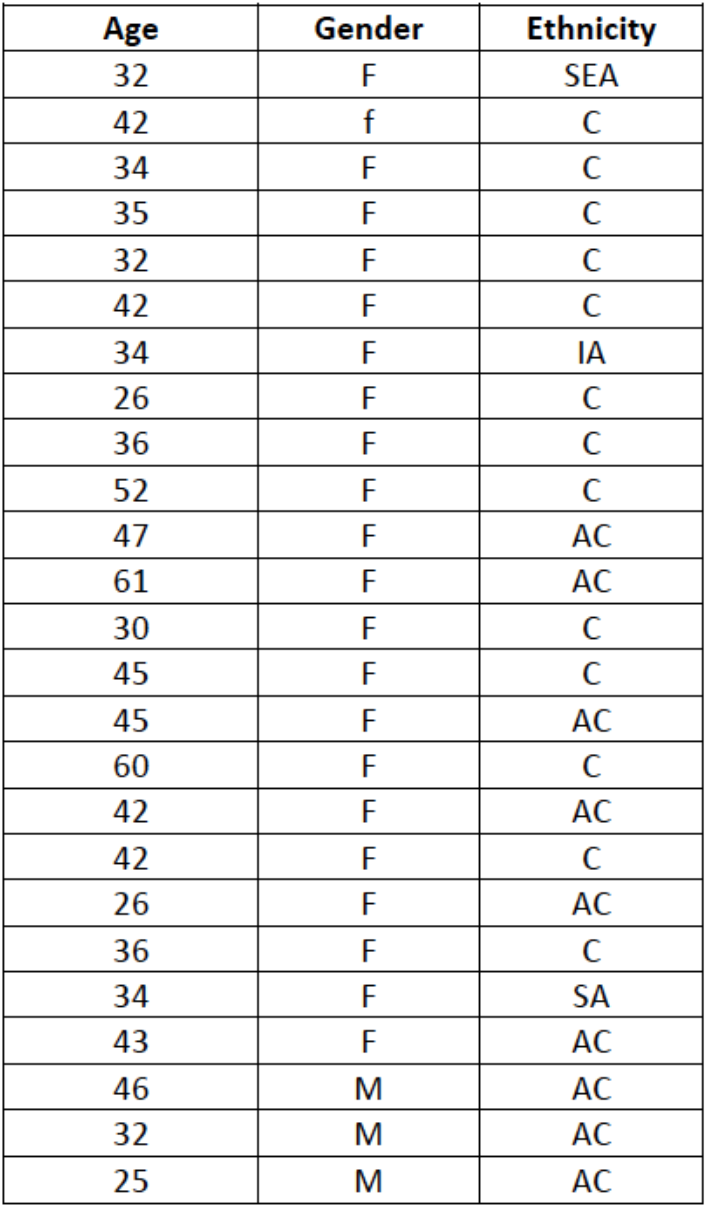
HCD demographic data used in Fig. 6. SEA = south east Asian, IA = Indian Asian.

| Age | Gender | Ethnicity |
| --- | --- | --- |
| 32 | F | SEA |
| 42 | f | C |
| 34 | F | C |
| 35 | F | C |
| 32 | F | C |
| 42 | F | C |
| 34 | F | IA |
| 26 | F | C |
| 36 | F | C |
| 52 | F | C |
| 47 | F | AC |
| 61 | F | AC |
| 30 | F | C |
| 45 | F | C |
| 45 | F | AC |
| 60 | F | C |
| 42 | F | AC |
| 42 | F | C |
| 26 | F | AC |
| 36 | F | C |
| 34 | F | SA |
| 43 | F | AC |
| 46 | M | AC |
| 32 | M | AC |
| 25 | M | AC |

**Table S3.** Demographic data of patients with other autoimmune diseases. GPA = granulomatosis with polyangiitis, PV = pemphigus vulgaris, UC = ulcerative colitis, AZA = azathioprine, MTX = methotrexate, MES = mesalazine.

| Age | Autoimmune disease | Sex | Medication |
| --- | --- | --- | --- |
| 34 | GPA | F | PRED, AZA |
| 49 | GPA | M | PRED |
| 63 | GPA | M | PRED, MTX |
| 68 | GPA | M | PRED, MTX |
| 62 | GPA | M | PRED, MTX |
| 34 | GPA | F | PRED, MTX |
| 42 | PV | M | PRED, MMF |
| 54 | PV | M | PRED, MMF |
| 39 | PV | F | PRED, MMF |
| 69 | PV | M | PRED, MMF |
| 82 | PV | M | PRED, MMF |
| 23 | PV | F | PRED, AZA |
| 24 | UC | F | MES |
| 28 | UC | F | PRED, AZA, MES |
| 64 | UC | F | MES |
| 23 | UC | M | PRED, AZA |
| 39 | UC | M | MES |
| 24 | UC | F | PRED, AZA, MES |
| 29 | UC | F | AZA, MES |
| 23 | UC | M | AZA, MES |

